# Ataxin-2 Coordinates Neuronal and Glial Pathways to Promote Neuroprotection

**DOI:** 10.1101/2025.11.24.690282

**Authors:** Denise Gastaldo, Steve Vito, Holly Black, Oded Foreman, Zhichang Yang, Victoria Pham, Ying Zhu, Meena Choi, Ana Jovičić

## Abstract

Ataxin-2 (ATXN2) is a genetic modifier of TDP-43 toxicity and a promising therapeutic target in amyotrophic lateral sclerosis (ALS). However, the mechanisms underlying its neuroprotective effects remain poorly understood. Here we show that ataxin-2 reduction confers neuroprotection by engaging adaptive metabolic programs in both neuronal and glial cells. Using global proteomic profiling in yeast and mouse TDP-43 models, we establish that pbp1/ataxin-2 (pbp1 is the yeast ataxin-2 ortholog) activates orthogonal stress-adaptive programs that enable alternative energy production and augment trophic support, rather than simply reversing TDP-43-induced damage. In neurons, ataxin-2 downregulation activates glycolysis and reductive glutamine carboxylation driven by IDH1, restoring ATP production independently of impaired mitochondria. In astrocytes, ataxin-2 downregulation upregulates cholesterol biosynthesis via HMGCS1, enhancing trophic support to neurons in a non-cell-autonomous manner. Full neuroprotection requires both mechanisms: neuronal survival is only completely rescued when ataxin-2 is reduced in the context of neuron-astrocyte co-cultures. Importantly, ataxin-2 downregulation protects against both TDP-43 gain-of-function and loss-of-function toxicity, broadening its therapeutic relevance. Collectively, our findings reveal novel mechanisms by which ataxin-2 orchestrates adaptive metabolic resilience and establish ataxin-2 as a regulator of stress-adaptive response in brain cells under the conditions of stress.

## Introduction

Amyotrophic Lateral Sclerosis (ALS) is a rapidly progressive and fatal neurodegenerative disease characterized by upper and lower motor neuron loss, with a typical survival of 2 to 5 years post-diagnosis. Existing ALS treatments are largely palliative, with modest efficacy at best, underscoring a significant unmet need for effective therapies. ALS is pathologically defined by the nuclear depletion and cytoplasmic accumulation of the RNA-binding protein TDP-43. Both of these pathological features are implicated in ALS pathogenesis, as evidenced by patient data and human genetics. Cytoplasmic TDP-43 aggregates are toxic to cells^1,2^, and variants in the TDP-43 prion-like domain that render it more aggregation-prone can cause ALS^3^. Conversely, nuclear TDP-43 loss results in dysregulation of mRNA polyadenylation^4^ and aberrant inclusion of cryptic exons^5,6,7^. This interplay between loss-of-function and gain-of-function mechanisms renders TDP-43 extremely difficult to target therapeutically. A promising strategy to bypass this challenge is to target genetic modifiers of TDP-43 that can confer cellular protection regardless of which TDP-43 pathological mechanism predominates.

Intermediate-length polyglutamine expansions in the range of 27–33 repeats in ataxin-2 (ATXN2) are associated with increased ALS risk across diverse human populations⁸⁻¹¹, establishing it as a common genetic risk factor for the disease. Ataxin-2 has been identified as a potent genetic modifier of TDP-43 toxicity⁸˒¹², and preclinical studies in yeast, fly, and mouse ALS models have demonstrated that reducing levels of wild-type ATXN2 homologs provides robust protection against TDP-43-associated cellular toxicity. These findings have positioned ataxin-2 downregulation as a promising therapeutic strategy for ALS, and a clinical trial of an *ATXN2*-targeting antisense oligonucleotide (ASO) was initiated. The clinical trial was discontinued without clinical benefit, possibly reflecting insufficient target engagement in the CNS, evidenced by only a modest ataxin-2 reduction observed in cerebrospinal fluid (CSF) [Ravits J, European Network for the Cure of Amyotrophic Lateral Sclerosis, 20th Meeting 19 June 2024]. Such results stand in stark contrast to preclinical studies, including the present work, where therapeutic rescue is typically predicated on the robust knockdown or genetic ablation of ataxin-2.

Despite this therapeutic interest, the cellular mechanism by which wild-type ataxin-2 downregulation confers neuroprotection has remained elusive. Here we show that ataxin-2 downregulation does not simply reverse the molecular damage caused by TDP-43. Instead, it activates adaptive metabolic programs that extend beyond a simple reversal of the TDP-43-induced damage, encompassing both cell-autonomous neuronal reprogramming and non-cell-autonomous glial adaptations. In neurons, we identified activation of ATP production through glycolysis and reductive glutamine carboxylation that compensate for impaired mitochondria. In astrocytes, we identify upregulation of cholesterol biosynthesis that enhances trophic support to neurons. Full neuroprotection requires both mechanisms: neuronal survival is completely rescued only when ataxin-2 is reduced in the context of neuron-astrocyte co-cultures. We establish this framework through proteomic profiling in yeast and mouse TDP-43 models, followed by functional validation in primary neuronal and neuron-astrocyte co-culture systems. We further show that ataxin-2 downregulation protects against both TDP-43 gain-of-function and loss-of-function toxicity, broadening its mechanistic and therapeutic relevance. Together, these findings reframe ataxin-2 as a regulator of stress-adaptive metabolic plasticity, identify isocitrate dehydrogenase (NADP(+)) 1 (IDH1) and 3-hydroxy-3-methylglutaryl-CoA synthase 1 (HMGCS1) as downstream effectors, and highlight the potential of targeting central nervous system metabolism in combating neurodegeneration.

## Results

### Pbp1 and ataxin-2 loss affect different cellular pathways than TDP-43 toxicity

Yeast ortholog of human *ATXN2* is *pbp1*^1^. The protective effects of *pbp1* deletion against TDP-43 expression toxicity in *Saccharomyces cerevisiae* have been well established and described previously^1^. This *S. cerevisiae* model is based on galactose-inducible overexpression of human TDP-43, which causes TDP-43 aggregation and cellular toxicity, recapitulating key aspects of TDP-43 proteinopathy in a heterologous system. To better understand the protective mechanisms at play, we conducted a comprehensive global proteomics analysis of *pbp1Δ* and wild-type yeast cultures during their exponential growth phase, eight hours following the induction of TDP-43 expression (Supplementary Table 1). Our initial observation revealed that TDP-43 expression in yeast cells results in an overall trend towards protein downregulation. Investigation of proteins significantly downregulated by TDP-43 revealed a group of mitochondrial proteins, including ATP synthase subunits, and ribosomal proteins (Fig. 1a). Deletion of *pbp1* under TDP-43 stress conditions led to changes in expression of mitochondrial proteins, both mitochondrial inner membrane proteins, tricarboxylic acid cycle and mitochondrial translation proteins. Interestingly, *pbp1Δ* triggered a metabolic shift characterized by the increased abundance of peroxisomal proteins, including fatty acid oxidation enzymes (Fig. 1b). Yeast cells typically resort to peroxisomal fatty acid oxidation when glucose availability becomes limited or when cellular energy demands exceed what can be met through glycolysis alone. This metabolic reprogramming suggests that *pbp1Δ* cells adapt to TDP-43-induced cellular stress by diversifying their energy production pathways. Metabolic reprogramming response was stress dependent as *pbp1* deletion in control GFP-expressing yeast resulted in different proteomic response (Supplementary Fig. 1, Supplementary Table 1). Our yeast findings point to two critical aspects of *pbp1Δ* protection against TDP-43: first, *pbp1* deletion does not rescue yeast by simply reversing the pathways affected by TDP-43 toxicity, but rather induces a distinct metabolic pathway that confers cellular resilience under conditions of stress (Fig. 1c). Second, given that neurons are not metabolically independent from their surrounding glia, this metabolic reprogramming highlights the need to consider non-cell-autonomous effects of ataxin-2 downregulation in the context of neuroprotection in mammalian cells.

**Figure 1.**
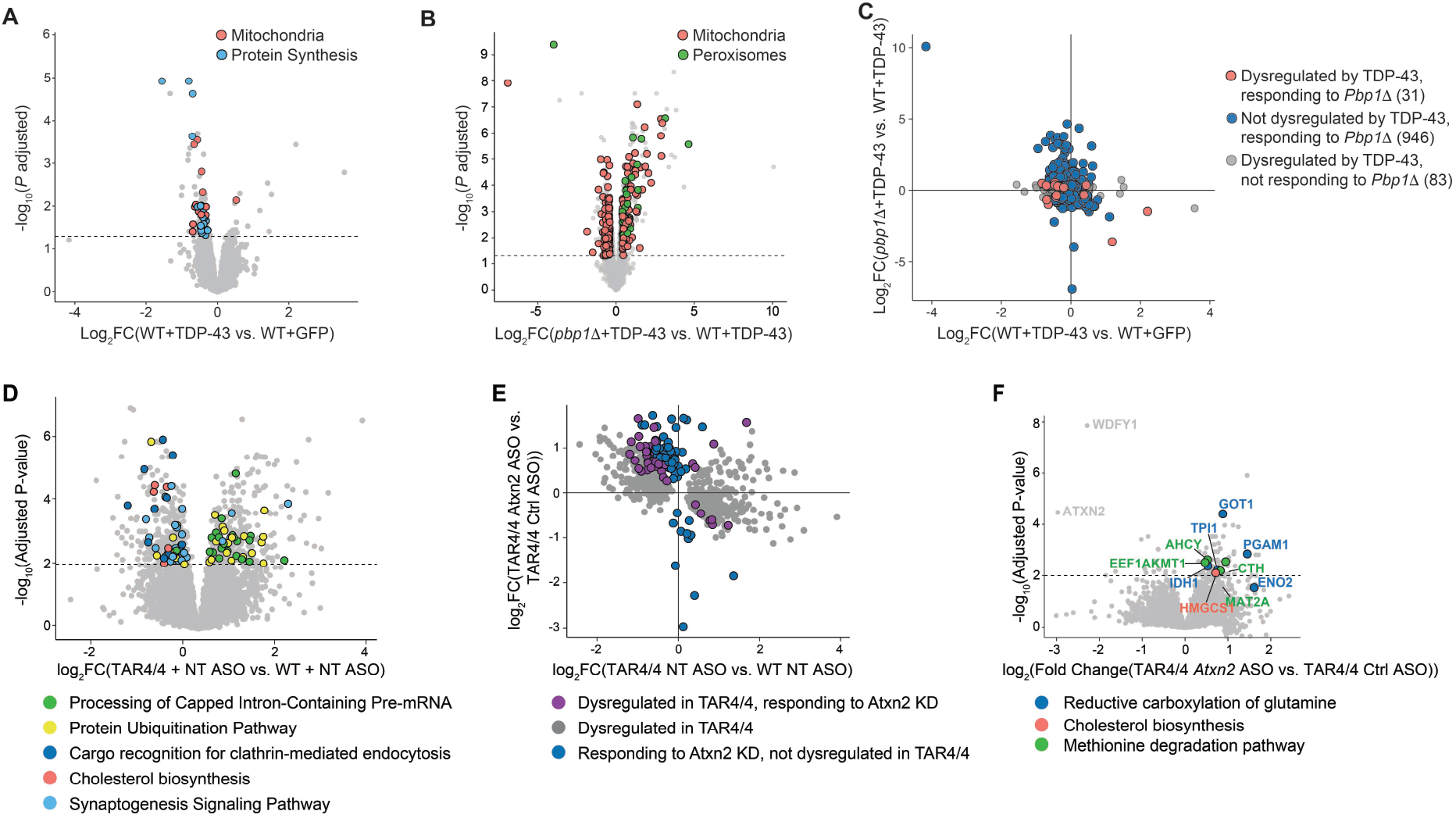
*Pbp1Δ* and ataxin-2 downregulation induce protective metabolic pathways upon TDP-43 expression. (A) Volcano plot showing differentially expressed proteins in wild-type yeast expressing TDP-43 versus control GFP (mass spectrometry; dashed horizontal line marks the adjusted *P* = 0.05). Data points are colored by Gene Ontology components identified using DAVID functional annotation tools. n=4 (WT + GFP), n=5 (WT + TDP-43). (B) Volcano plot showing differentially expressed proteins in *pbp1Δ* yeast expressing TDP-43 versus control GFP (mass spectrometry; dashed horizontal line marks the adjusted P = 0.05). Data points are colored by Gene Ontology components identified using DAVID functional annotation tools. n=3 (*pbp1Δ* + GFP), n=4 (*pbp1Δ* + TDP-43). (C) Comparing the effects of *pbp1Δ* on the *y* axis to the effects of TDP-43 expression on *x* axis. Proteins with adjusted *P* < 0.05 are shown. The number of significantly changed proteins in each category is indicated in parentheses. (D) Volcano plot of differentially expressed proteins in layer 5 motor cortex of TAR4/4 versus wild-type mice (mass spectrometry; *P* adjusted < 0.01). n=4 male mice per group. Proteins colored by IPA pathway annotation. (E) Scatter plot comparing disease-related protein changes (*x* axis: TAR4/4 vs. wild-type, both control ASO treatment) with ataxin-2 downregulation effects (*y* axis: TAR4/4, *Atxn2* ASO vs. TAR4/4, control ASO) in layer V cortex. Significantly altered proteins shown (*P* adjusted < 0.01, mass spectrometry). n=4 male mice per group. (F) Volcano plot of differentially expressed proteins in layer V motor cortex of TAR4/4 mice *Atxn2* ASO vs. control ASO treatment (mass spectrometry, *P* adjusted < 0.01). Proteins are colored by pathway. n=4 male mice per group.

### Ataxin-2 downregulation activates orthogonal metabolic pathways in ALS mouse brain

To further define the cellular pathways activated by ataxin-2 downregulation, we performed comprehensive proteomic profiling in mouse brain tissue of TAR4/4 ALS model mice treated with an antisense oligonucleotide (ASO) against ataxin-2. TAR4/4 mice are a recognized standard for studying ALS-associated neurodegeneration, as they consistently develop a severe, lethal phenotype characterized by TDP-43 overexpression and motor decline^2^. Additionally, they have been previously validated as a responsive model for Atxn2-targeting ASO injection, thus providing a suitable model to study the mechanism of the Atxn2-TDP-43 genetic interaction^3^. Analogous to pbp1 in yeast, ataxin-2 downregulation, rescues neurodegeneration and extends the lifespan in mouse TAR4/4 ALS model (Supplementary Figure 2A and B)^3^. Due to the particular relevance of cortical layer V to ALS, and presence of neurodegeneration (Supplementary Figure 2C) with limited extent of cell death in this region (Supplementary Figure 2D) providing an opportunity to interrogate pre-apoptotic cellular processes, we performed spatially-resolved proteomic analysis by microdissecting cortical layer V and analyzed these samples using a highly sensitive method^4,5^. As layer V contains a heterogeneous population of cell types, including neurons, glia, and vascular cells, rather than a pure neuronal population, we performed cell-type marker analysis in parallel to inform interpretation of the proteomic changes detected (see below).

Proteomic analysis (Supplementary Table 2) revealed strong dysregulation of protein expression in TAR4/4 layer V. Ingenuity Pathway Analysis (IPA)^6^ identified protein ubiquitination and processing of capped intron-containing pre-mRNAs among the top upregulated pathways, while synaptogenesis and cholesterol biosynthesis pathway emerged among the most prominently downregulated pathways (Figure 1D, Supplementary Table 2). When examining cellular response to ataxin-2 downregulation in TAR4/4 mice we detected a partial reversal of TAR4/4-associated dysregulation, but also orthogonal protein changes not observed in this disease model (Figure 1E). This is a pattern that mirrors our yeast findings, where pbp1 loss effects were mostly distinct from TDP-43 toxicity. In TAR4/4 mice, ataxin-2 silencing mostly led to protein upregulation. Proteins upregulated in response to ataxin-2 loss in TAR4/4 include enzymes involved in glycolysis (TPI1, PGAM1, ENO2), reductive glutamine carboxylation pathway (GOT1, IDH1), HMGCS1 (a critical enzyme in the mevalonate cholesterol synthesis pathway), and components of methionine degradation pathway (CTH, AHCY, EEF1AKMT1, MAT2A) (Figure 1F). The upregulated enolase was the neuron-specific isoform ENO2, suggesting that upregulation of glycolysis occurs specifically in neurons and represents a cell-autonomous protective mechanism. The upregulation of cholesterol and methionine degradation pathways is particularly intriguing, as these processes occur predominantly in astrocytes within the brain^7,8^. Given that layer V microdissections necessarily contain a mixed population of cell types beyond neurons and astrocytes, we performed cell-type marker analysis to evaluate whether the observed proteomic changes could be explained by shifts in cell-type composition following Atxn2 ASO treatment (Supplementary Figure 3A, B). This analysis did not reveal substantial block-level differences in cell-type marker abundance between TAR4/4 mice treated with control ASO versus Atxn2 ASO, suggesting that the proteomic changes we observed are unlikely to be primarily driven by large shifts in the cellular composition of the tissue. This observation supports the interpretation that ataxin-2 downregulation may confer neuroprotection through modulation of astroglial protein expression, revealing a potential non-cell-autonomous protective mechanism that extends beyond direct neuronal effects. Finally, only 24 proteins responded to ataxin-2 downregulation in wild-type mice (Supplementary Figure 4, Supplementary Table 2) and all of them were downregulated, indicating that the metabolic effects of ataxin-2 loss are stress-dependent.

### Ataxin-2 loss protects neurons via both neuronal cell-autonomous and astroglial non cell-autonomous manner

To interrogate the pathways identified through proteomics analyses, we established an *in vitro* model of TDP-43 toxicity in primary cortical neurons, using NeuN+ cell counts as a quantitative endpoint of neuronal survival. TDP-43 overexpression did not lead to detectable TDP-43 aggregation in these cells. Cortical upper motor neurons are among the cardinal cellular targets of neurodegeneration in ALS, and the primary motor cortex correspondingly exhibits regional glucose hypometabolism, as evidenced by F-FDG-PET imaging^9^. TDP-43 overexpression in cortical primary neurons induced substantial toxicity and cell death (Figures 2A and 2B), accompanied by severe mitochondrial dysfunction marked by impaired oxygen consumption rate (OCR) and diminished ATP production (Figures 2C and 2D), consistent with prior findings^10–12^. Although treatment of primary neurons with *Atxn2*-targeting ASO strongly reduced *Atxn2* levels (Supplementary Figure 5A), it produced only a modest, non-significant improvement in neuronal survival (Figures 2A and 2B), did not improve OCR (Figure 2C) but increased total ATP production (rescue effect of ∼50%) (Figure 2D). However, when neurons were co-cultured with astrocytes, *Atxn2* ASO treatment of the cocultures achieved rescue of neuronal viability (Figure 2E, F, Supplementary Figure 5B), demonstrating that ataxin-2 downregulation confers protection through both cell-autonomous neuronal mechanisms and non-cell-autonomous astroglial pathways.

**Figure 2.**
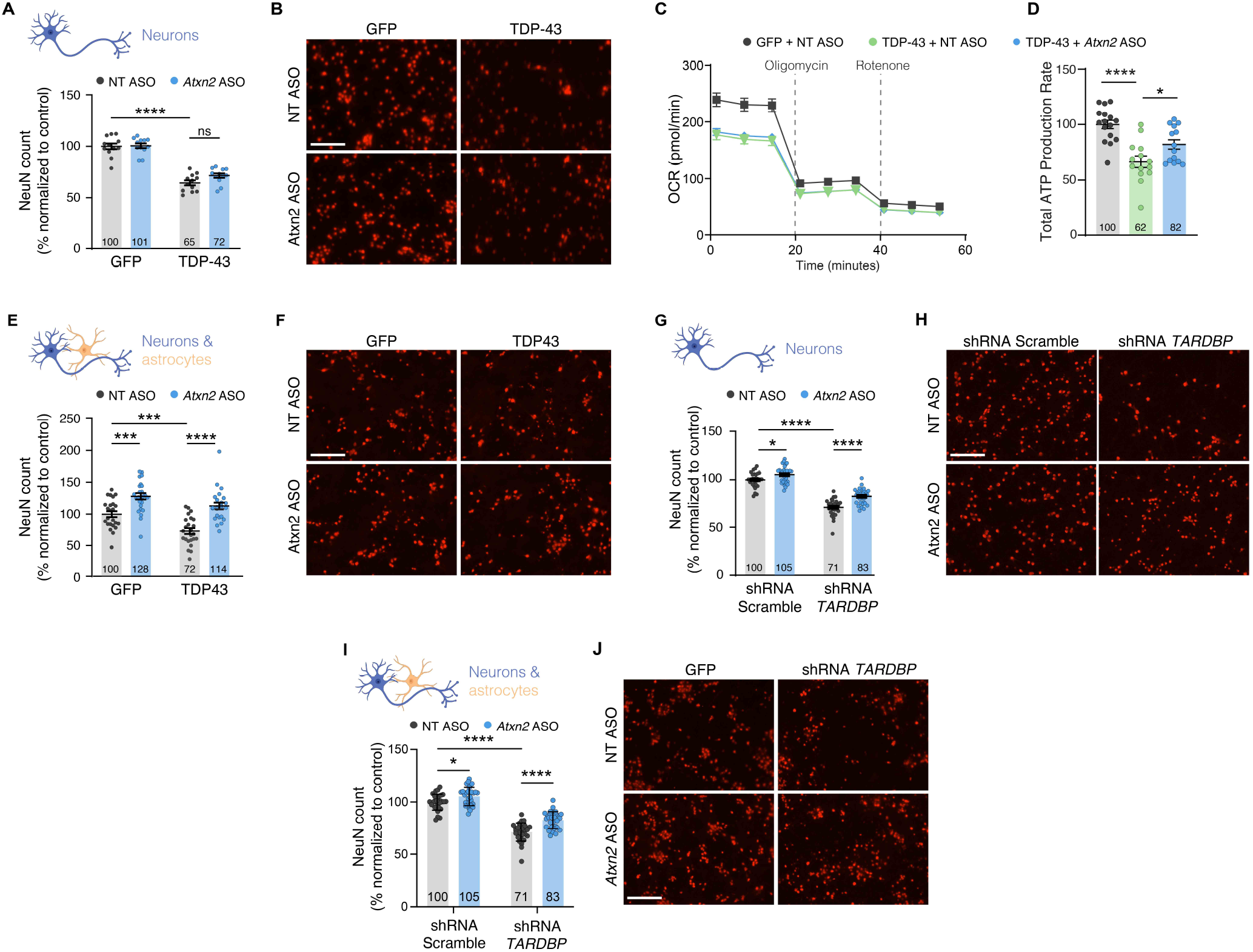
Ataxin-2 downregulation promotes neuronal health *in vitro* via cell-autonomous and non cell-autonomous effects. (A) Quantification of primary neuron survival in control (GFP) or TDP-43 overexpressing conditions, treated with NT or *Atxn2* ASO. Data are normalized to GFP with NT ASO condition. (B) Representative images of NeuN-labeled neurons quantified in (A). (C) Oxygen Consumption Rate (OCR, pmol min^-1^) over time in GFP or TDP-43 overexpressing neurons treated with NT or *Atxn2* ASO. (D) Total ATP production rates in primary neurons overexpressing TDP-43 treated with NT or *Atxn2* ASO, compared to GFP control neurons treated with NT ASO. ATP production rates were normalized to cell number and expressed as relative to GFP with NT ASO condition. (E) Quantification of neuronal survival in neuron and astrocyte co-cultures, in GFP or TDP-43 overexpressing conditions, treated with NT or *Atxn2* ASO. Data are normalized to GFP with NT ASO condition. (F) Representative images of NeuN-labeled neurons co-cultured with astrocytes quantified in (E). (G) Primary neuron survival in control (shRNA Scramble) or TDP-43 loss (shRNA *TARDBP*) conditions, treated with NT or *Atxn2* ASO. (H) Representative images of NeuN-labeled neurons quantified in (G). (I) Quantification of neuronal survival in neuron and astrocyte co-cultures, in control (shRNA Scramble) or TDP-43 loss (shRNA *TARDBP*) conditions, treated with NT or *Atxn2* ASO. Data are normalized to control with NT ASO condition. (J) Representative images of NeuN-labeled neurons co-cultured with astrocytes quantified in (I). All data are means ± SEM; statistical significance was assessed by two-way ANOVA with Tukey’s multiple comparison test (A) or Šidák’s multiple comparison test (E, G, I), or one-way ANOVA with Šidák’s multiple comparison test (D), with * *P* adjusted < 0.05, ** *P* adjusted < 0.01, *** *P* adjusted < 0.001, and **** *P* adjusted < 0.0001, ns is nonsignificant; scale bars = 200 µm.

### Ataxin-2 downregulation protects against TDP-43 loss of function toxicity

We observed that ataxin-2 knockdown improved survival broadly, extending survival not only in TDP-43 overexpression but also in control GFP-expressing cells (Figure 2). We further confirmed that GFP expression used as a control in our experiments did not impair neuronal viability and that *Atxn2* ASO exerts the same prosurvival effect in naïve cultures (Supplementary Figure 6A and B). Because even naïve cultures are subject to the baseline stress of *in vitro* conditions, this finding indicates that ataxin-2 downregulation confers pro-survival effects that may be efficacious beyond TDP-43 overexpression toxicity. We next tested whether this protection extends to TDP-43 loss-of-function toxicity. Ataxin-2 downregulation partially rescued neuronal survival caused by TDP-43 loss, and the effect was stronger in neuron-astrocyte cocultures than neuronal cultures alone (Figures 2G-J). This finding is particularly relevant given that nuclear TDP-43 loss is a defining feature of ALS pathology^13^, and indicates that ataxin-2 downregulation can confer protection against both arms of TDP-43 dysfunction, a key advantage in a disease where gain- and loss-of-function mechanisms coexist.

### Ataxin-2 downregulation rescues neuronal ATP production *in vitro*

Our proteomic profiling of layer 5 in TAR4/4 mice indicated that ataxin-2 downregulation leads to an upregulation of enzymes involved in glycolysis and reductive glutamine carboxylation pathways. Increased glycolysis can provide additional energy to cells with impaired mitochondria, while activation of reductive glutamine carboxylation can further support glycolysis through NADH shuttling^14,15^. Oxygen consumption rate (OCR) and extracellular acidification rate (ECAR), reflecting mitochondrial respiration and proton production from glycolysis respectively, were both impaired in neurons overexpressing TDP-43 (Figures 2C, 3A and 3B). Ataxin-2 loss had no effect on OCR, but rescued glycoATP production rate (∼41% rescue) and ECAR, demonstrating that ataxin-2 reduction stimulates neurons to produce more ATP through glycolysis (Figure 3A) and not by repairing mitochondrial dysfunction. To demonstrate that these metabolic effects are supported by reductive glutamine carboxylation pathway we silenced the key enzyme of the pathway, IDH1 (Supplementary Figure 6A). This completely abolished the rescue effect of ataxin-2 downregulation on ATP production and ECAR (Figures 3C and D). To further test whether activated reductive glutamine carboxylation is able to enhance ATP synthesis and survival in neurons *in vitro*, we overexpressed IDH1 (Supplementary Figure 6B). IDH1 overexpression alone was sufficient to increase ATP production through glycolysis (∼46% rescue) (Figure 3E and F) and neuronal survival (Figure 3G and Supplementary Figure 6C) validating our hypothesis. Remarkably, IDH1 overexpression provided greater neuronal survival benefit than ataxin-2 knockdown in primary neuronal cultures, possibly reflecting the higher IDH1 protein levels (we were not able to detect IDH1 protein using western blotting with commercially available antibodies). Collectively, this data indicates that ataxin-2 downregulation *in vitro* protects against TDP-43-induced mitochondrial dysfunction, by activating glycolysis and reductive glutamine carboxylation, thus enabling neurons to synthesize ATP (Figure 3I). In addition, they suggest IDH1 is a particularly potent downstream effector of this pathway.

**Figure 3.**
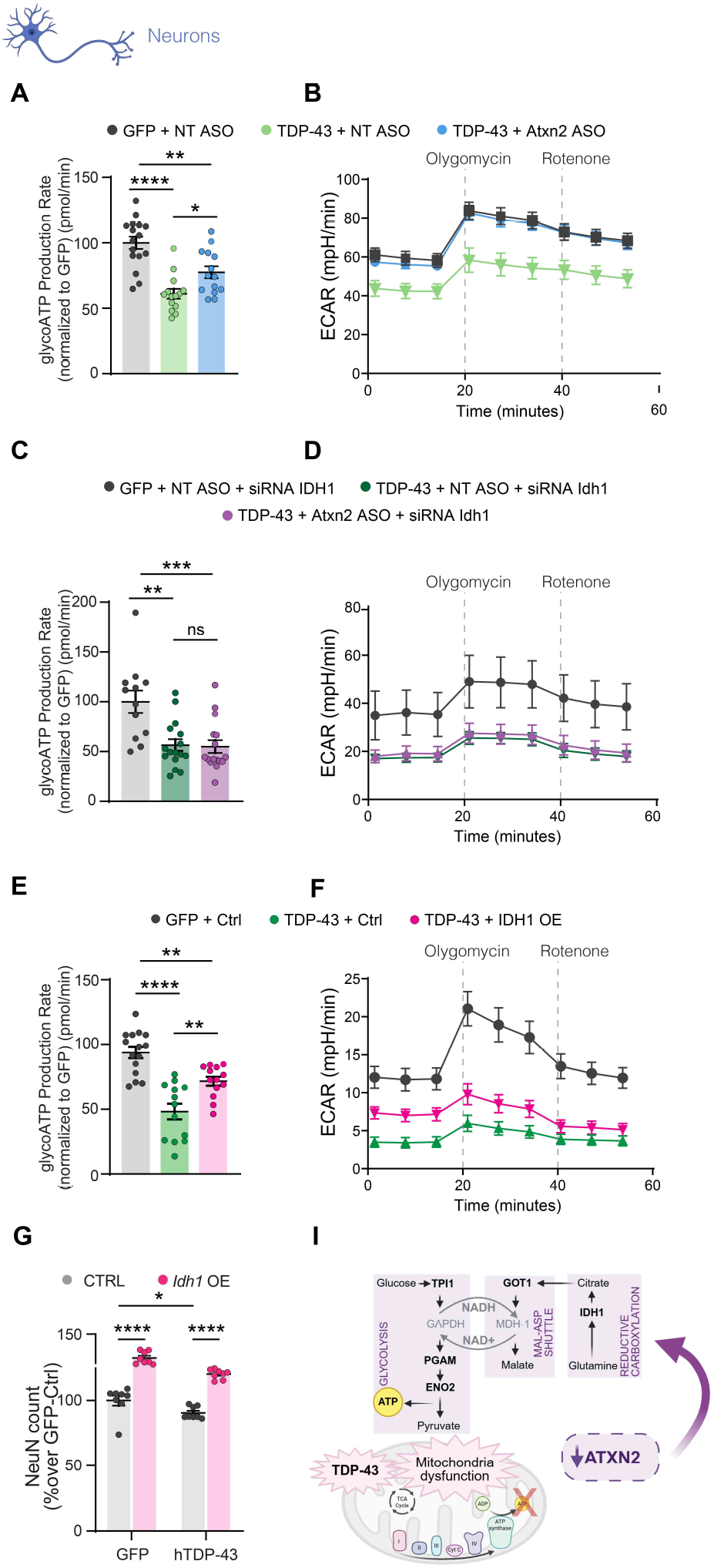
Ataxin-2 reduction in neurons enhances ATP synthesis via reductive glutamine carboxylation and glycolysis, promoting their survival. (A, B), Glycolytic ATP (glycoATP) production rates (pmol/min) (A) and Extracellular Acidification Rate (ECAR) (B) of GFP- or TDP-43 overexpressing neurons, treated with NT or *Atxn2* ASO. (C, D) GlycoATP (C) and ECAR (D) of GFP- or TDP-43 overexpressing neurons, treated with NT or *Atxn2*ASO, and *Idh1* siRNA. (E, F) GlycoATP (E) and ECAR (F) of GFP- or TDP-43 overexpressing neurons, treated with NT or *Atxn2* ASO, and overexpressing GFP (Ctrl) or IDH1. (G) Effect of IDH1 overexpression on TDP-43 toxicity, as measured by NeuN counts. Data are presented as normalized over GFP controls. (I) Simplified scheme of reductive glutamine carboxylation upon mitochondria dysfunction. Enzymes involved in the pathway and found upregulated upon *Atxn2* ASO in the mouse brain are highlighted in bold. All data are means ± SEM; statistical significance was assessed by one-way ANOVA with Tukey’s multiple comparison test (in A, C, E) or Two-way ANOVA followed by Šidák’s multiple comparison test (G), with * *P* adjusted < 0.05, ** *P* adjusted < 0.01, *** *P* adjusted < 0.001, and **** *P* adjusted < 0.0001,

### Ataxin-2 loss boosts cholesterol production in astrocytes *in vitro*

Given our proteomics results and known role of TDP-43 in regulating cholesterol synthesis^16^, we hypothesized that the astroglial protection mechanism may at least in part be mediated through increased cholesterol synthesis. First, we verified that the mass spectrometry data were reproducible *in vitro*. Western blot performed in neuron-astrocyte co-culture experiments confirmed a reduction of HMGCS1 when neurons are overexpressing TDP-43, and HMGCS1 levels were partially restored upon ataxin-2 knockdown (Figures 4A and B). Next, we transduced astrocytes with a lentivirus expressing HMGCS1 and then used these astrocytes for co-culture and neuronal survival assay (Figures 4C, 4D, Supplementary Figure 7A). Overexpressing HMGCS1 in astrocytes was protective against TDP-43 toxicity in neurons. To further validate that ataxin-2 knockdown *in vitro* is protective through this pathway, we combined *Atxn2* ASO treatment with an siRNA targeting *Hmgcs1* (Supplementary Figures 7B, Figures 4E and F). Confirming our hypothesis, knock-down of HMGCS1 in co-cultures abolished the protection given by the ataxin-2 knockdown in astrocytes. To rule out a possible contribution of cholesterol biosynthesis in neurons, we overexpressed HMGCS1 in primary neuronal cultures and we did not observe any rescue of survival (Supplementary Figures 7C and D), giving additional evidence of the non-cell autonomous effect of ataxin-2 through cholesterol synthesis in astrocytes. Finally, impeding cholesterol release through downregulation of Abca1, key cholesterol efflux transporter^17^, abolished prosurvival effects of *Atxn2* ASO (Figures 4G and H). Together, these data indicate that HMGCS1, and its downstream effector ABCA1, are necessary and sufficient to mediate the astrocyte-derived cholesterol pathway that supports neuronal viability in this co-culture model (Figure 4I). This non-cell-autonomous mechanism acts in parallel with the neuron-autonomous IDH1-dependent pathway described above (Figs. 3), indicating that Atxn2 downregulation engages at least two distinct, convergent mechanisms of neuroprotection rather than a single unifying pathway.

**Figure 4.**
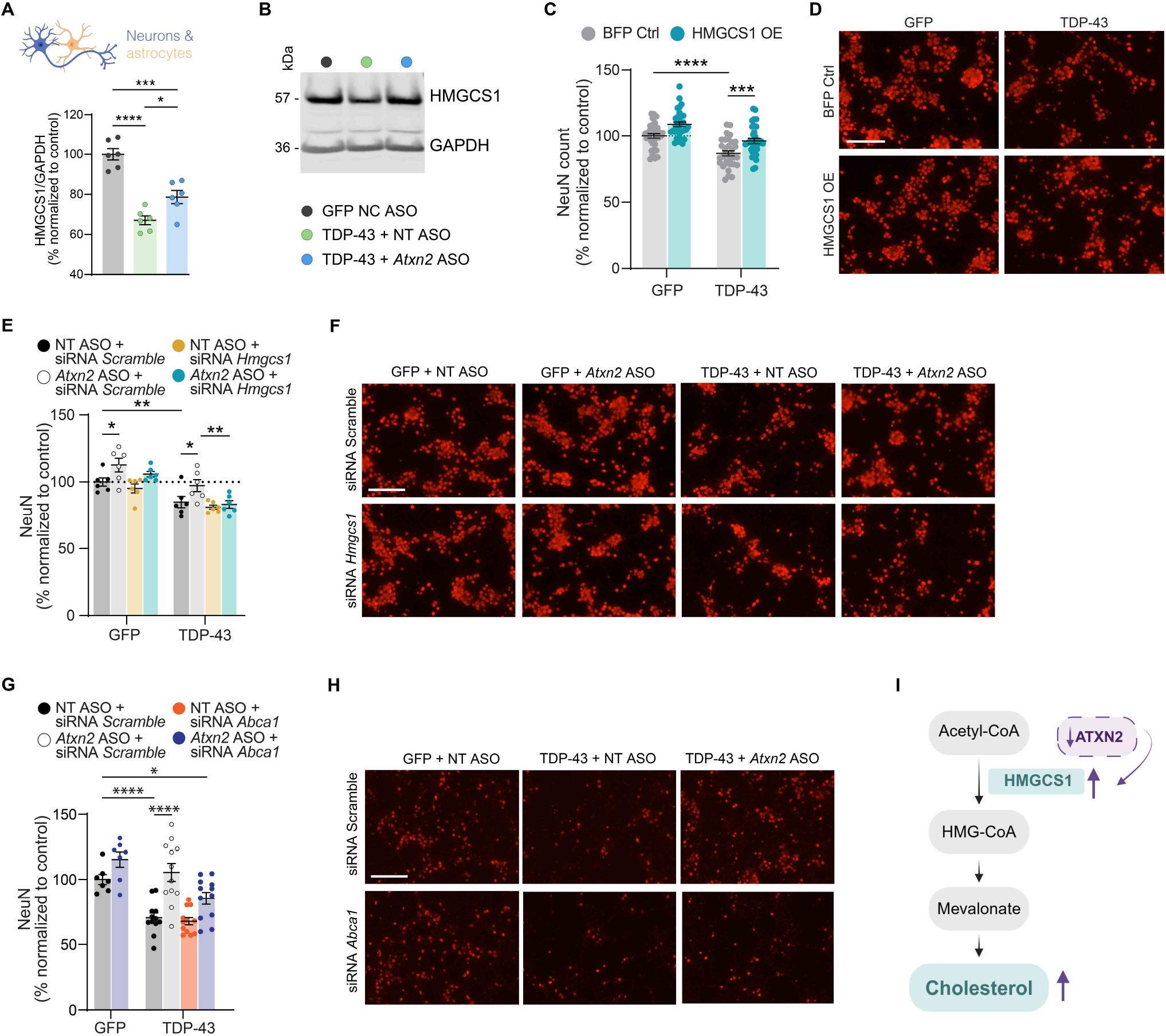
Ataxin-2 depletion in astrocytes enhances cholesterol biosynthesis, supporting neuronal survival. (A) Western blot analysis of HMGCS1 protein levels in astrocytes co-cultured with control GFP or TDP-43 overexpressing neurons, treated with NT or *Atxn2* ASO. Quantification shows signal intensity relative to GAPDH, normalized to GFP + NT ASO condition. (B) Representative cropped western blots of proteins analyzed in (A). (C) NeuN-positive cell counts in co-cultures of TDP-43- or GFP-expressing neurons, plated on astrocytes overexpressing HMGCS1 or BFP control. Data are normalized to BFP control conditions. (D) Representative images of primary neurons overexpressing TDP-43 plated on astrocytes that overexpress HMGCS1 and stained with NeuN (scale bar = 200 µm). (E) NeuN-positive cell counts in TDP-43 overexpressing or GFP-expressing neurons co-cultured with astrocytes following *Atxn2* ASO and *Hmgcs1* siRNA treatment. Data are normalized to GFP + NT ASO + scrambled (Scr) siRNA condition. (F) Representative images of NeuN staining in co-cultures from *Hmgcs1* siRNA experiment quantified in (E) (scale bar = 200 um). (G) NeuN-positive cell counts in TDP-43 overexpressing or GFP-expressing neurons co-cultured with astrocytes following *Atxn2* ASO and *Abca1* siRNA treatment. Data are normalized to GFP + NT ASO + scrambled (Scr) siRNA condition. (H) Representative images of NeuN staining in co-cultures from *Abca1* siRNA experiment quantified in (G) (scale bar = 200 um). (I) Simplified schematic of the mevalonate pathway for cholesterol biosynthesis. HMGCS1, which is upregulated upon ataxin-2 knockdown in proteomic analysis, is the key enzyme catalyzing the first reaction of the pathway. Purple arrows indicate the effect of ataxin-2 downregulation on HMGCS1 expression and downstream effects on cholesterol biosynthesis. All data are means ± SEM; statistical significance was assessed by one-way ANOVA followed by Tukey’s multiple comparison test (in A) or Two-way ANOVA followed by Šidák’s multiple comparison test (C, G) with * *P* adjusted < 0.05, ** *P* adjusted < 0.01, *** *P* adjusted < 0.001, and **** *P* adjusted < 0.0001, and two-way ANOVA with Fisher’s LSD multiple comparison test (E) * *P* < 0.05, ** *P* < 0.01

## Discussion

This study uncovers a metabolism-centered framework for how pbp1/ataxin-2 downregulation counteracts TDP-43 toxicity across evolutionarily distant systems. Rather than simply reversing the molecular perturbations caused by TDP-43, pbp1/ataxin-2 reduction initiates adaptive metabolic response that allows cells to sustain energy production and trophic support under stress. In both yeast and neural cells, pbp1/ataxin-2 reduction diversifies energy production away from compromised mitochondria toward alternative pathways. In yeast, *pbp1* deletion activates peroxisomal fatty acid oxidation, while in neurons, *Atxn2* silencing activates glycolysis and reductive glutamine carboxylation. In addition, in astrocytes, *Atxn2* silencing enhances cholesterol biosynthesis to offer trophic support which bolsters neuronal survival in a non–cell-autonomous manner. Together, these data identify ataxin-2 as a regulator of stress-adaptive metabolic plasticity in cells exposed to TDP-43 toxicity. The difference between pathways perturbed by TDP-43 and those engaged by ataxin-2 loss (observed in both yeast and mouse) argues that ataxin-2 reduction does not merely correct TDP-43–induced molecular defects but reprograms cellular state toward stress resilience. Previously, polyQ expansions in ataxin-2 have been linked to aberrant sequestration of TDP-43, with consequent perturbation of the spatial localization and local translation of axonal mRNAs^18^. In addition, prior studies showed that ataxin-2 functions as a crucial mediator in RNP granules formation, and highlighted that targeted reduction in ataxin-2-mediated RNP-granule formation through mutation of its specific domains can ameliorate neurodegeneration, including FUS and C9orf72 ALS models^19,20^. These findings may suggest that ataxin-2 absence or reduction could alter stress granule formation or content in response to cellular stress and prevent the functional sequestration and translation of specific mRNA involved in glycolysis, reductive carboxylation, and cholesterol biosynthesis, thus promoting survival. However, this molecular mechanism needs to be further investigated.

The cross-species consistency, from peroxisomal compensation in yeast to glycolysis/reductive flux and sterol pathways in mammals, argues that the metabolic regulation by ataxin-2 is phylogenetically conserved, and functionally relevant under stress.

Energy failure and lipid dysregulation are increasingly recognized as common features in ALS. Our *in vitro* data show that *Atxn2* silencing restores ATP via glycolysis/reductive glutamine carboxylation and increases cholesterol supply from astrocytes, suggesting that ataxin-2 is mechanistically linked to two recurrent vulnerabilities in ALS: neuronal bioenergetics and glia-mediated metabolic support. Multiple studies have shown through PET and autoradiography imaging that cerebral tissue shows glucose hypo metabolism^21^. Importantly, these changes are detected before cell death and are associated with disease progression^22^. In addition recent studies directly linked TDP-43 mislocalization with impaired glycolysis through its direct link with hexokinase 1, and demonstrate that glycolytic restoration is a potential therapeutic strategy in ALS (Barone et al., 2026). Ataxin-2 downregulation could directly alleviate glucose hypometabolism in affected neurons, correct the energetic crisis in ALS and provide resilience to degenerating motor neurons. Our *in vitro* studies highlight IDH1 as a potential pharmacologic target to achieve energetic resilience in neurons. This raises the possibility that IDH1-directed strategies could serve as a complementary or alternative therapeutic avenue, particularly in contexts where ataxin-2 lowering is difficult to achieve at a sufficient threshold in relevant CNS regions.

Furthermore, we observed that ataxin-2 downregulation engages both cell-autonomous neuronal adaptation, as well as non cell-autonomous astroglial adaptation. TDP-43 impairs cholesterol synthesis by inhibiting SREBP2 activity and downregulating its targets, including HMGCS1^16^. Loss- and gain-of-function experiments demonstrate that HMGCS1, together with ABCA1-mediated cholesterol efflux, is both necessary and sufficient for this astrocytic protective pathway; however, this pathway acts alongside, rather than in place of, the neuron-intrinsic IDH1-dependent mechanism, indicating that Atxn2-mediated neuroprotection reflects the convergence of at least two distinct pathways rather than a single dominant mechanism. In addition, while patient data suggest that higher cholesterol is associated with increased risk of developing ALS^23^, after ALS diagnosis, higher cholesterol levels correlates with a lower risk of mortality^24^. Therefore, upregulating astroglial cholesterol transport to neurons after ALS diagnosis could provide the needed support to sick neurons. Importantly in wild-type mice, lowering ataxin-2 levels resulted in very few protein expression changes. The pathways we identified as neuroprotective were not upregulated in wild-type mice treated with *Atxn2* ASO. These observations suggest a potentially highly favorable safety profile of *Atxn2* silencing strategies in the clinic, as the intervention would impact stressed cells while leaving healthy cells largely unaffected. Exactly how ataxin-2 senses stress signals in diseased neurons remains unknown. Speculatively, stress-associated signals, such as metabolic imbalance, energy crisis or elevated reactive oxygen species (ROS), could be sensed by ataxin-2, thereby modulating its role in stress-responsive pathways, defining a promising field of research for future studies.

Notably, our proteomic studies were performed on a pre-apoptotic region in the mouse brain and therefore uncovered protective responses that can extend neuronal survival. The *in vitro* studies showed that manipulating ataxin-2 levels extends neuronal life span against both TDP-43 overexpression and TDP-43 downregulation, significantly expanding the relevance to ALS. In addition, our results nominate downstream targets (IDH1 and HMGCS1) that could complement or substitute for ataxin-2 lowering. In neurons, boosting IDH1-dependent reductive glutamine carboxylation or stabilizing cytosolic NADPH/NADH shuttles could preserve ATP under respiratory stress. In astrocytes, enhancing cholesterol biosynthesis and export could augment synaptic and membrane maintenance in degenerating circuits. These strategies could be particularly valuable if ataxin-2 suppression alone is insufficient or not uniformly achieved in relevant cell types. More broadly, our findings encourage a shift toward “metabolic resilience” therapeutics in ALS, addressing both neuronal bioenergetic failure and glial trophic deficits.

The recent early-phase trial of an *ATXN2* ASO reported modest CSF knockdown of ataxin-2 levels and did not show clinical benefit over the observation window (Clinical Trial NCT04494256; Ravits J, European Network for the Cure of Amyotrophic Lateral Sclerosis, 20th Meeting 19 June 2024). In our experiments, we achieved ataxin-2 downregulation greater than 90%. While future studies will help understand the relationship between central nervous system (CNS) and CSF ataxin-2 protein levels, interpretation of our results may indicate that therapeutic efficacy may require achieving a strong threshold of ataxin-2 lowering: sufficient ataxin-2 lowering in all relevant neurons to activate reductive glutamine carboxylation and in surrounding astrocytes to augment cholesterol biosynthesis. Importantly, understanding the mechanisms of neuroprotection may enable biomarker selection for future clinical trials to better understand target engagement in the clinic.

A limitation of this study is that the causal contribution of specific metabolic pathways, glycolysis/reductive glutamine carboxylation in neurons and cholesterol biosynthesis in astrocytes, to survival benefit *in vivo* has not been directly tested. Our *in vitro* manipulations of IDH1 and HMGCS1/ABCA1 support a role for these pathways in ataxin-2-mediated neuroprotection, but whether they are individually necessary or sufficient for the survival benefit observed in TAR4/4 mice remains to be established. Future studies will be needed to formally test the necessity and sufficiency of these metabolic pathways in vivo.

A further limitation is that our functional dissection of these metabolic pathways was performed exclusively in TDP-43-based models. Because TDP-43 dysfunction is common to the large majority of ALS cases, including C9orf72 and sporadic ALS, we anticipate that the metabolic vulnerabilities described here may extend beyond TDP-43-driven disease. However, we have not tested whether ataxin-2 downregulation similarly engages glycolysis/reductive glutamine carboxylation or astrocytic cholesterol biosynthesis in SOD1- or FUS-based models, which do not exhibit TDP-43 pathology, or in other models of familial ALS (for example C9orf72-based models) or sporadic ALS. Whether ataxin-2 lowering is protective in these models at all, and if so through the same or distinct mechanisms, remains to be tested.

A related consideration concerns the form of ataxin-2 studied here. Throughout this work, we reduced levels of wild-type pbp1/Atxn2 in yeast and mouse models. We did not directly test polyQ-expanded, ALS-associated ATXN2 alleles. Intermediate CAG repeat expansions in ATXN2 are a well-established genetic risk factor for ALS, but the functional consequences of these expansions are not fully understood, and it remains possible that expanded ATXN2 acts, at least in part, through a neomorphic mechanism. Our mechanistic analysis therefore addresses the stress-adaptive role of wild-type ataxin-2, and does not establish whether the same glycolytic, reductive carboxylation, and cholesterol biosynthesis programs are engaged in the context of polyQ-expanded ATXN2. We propose the metabolic programs identified here as candidate pathways that may be activated by ATXN2-lowering therapeutic strategies, which reduce total ATXN2 protein regardless of repeat length, rather than as an account of the primary pathogenic mechanism of polyQ-expanded ATXN2 in ALS patients.

In summary, ataxin-2 downregulation engages an adaptive metabolic program across neurons and astrocytes that protects cells against TDP-43–induced stress. Our findings suggest that by enabling neurons to generate ATP independently of damaged mitochondria and by increasing astrocytic cholesterol support, ataxin-2 targeting may address two central and convergent liabilities in ALS. These insights provide mechanistic rationale for refining ataxin-2-directed therapies, for developing complementary metabolic interventions, and for deploying aligned biomarkers to guide and de-risk clinical translation.

## Material and methods

### Yeast strains, media and plasmids

Yeast cells were grown in rich medium (YPD) or in synthetic media lacking uracil and containing 2% glucose (SD-Ura) or galactose (Sgal-Ura). TDP-43 and GFP yeast expression plasmids were prepared by VectorBuilder Inc, (Chicago, IL USA) by inserting *TARDBP* or *GFP* coding sequences into YEp plasmids with GAL promoter. All constructs were verified by DNA sequencing. YEp-GAP-TDP43 or YEp-GAL-GFP constructs were transformed into the BY4741 strain. Strains were manipulated and media prepared using standard techniques. We used the PEG/lithium acetate method to transform yeast with plasmid DNA. To induce TDP-43 or GFP expression for proteomics experiments, yeast cells were grown overnight at 30 °C in liquid Sglu-Ura medium until they reached log or mid-log phase. Cultures were then diluted to OD600 0.5 and grown in Sgal-Ura media at 30 °C for 8 hours.

### Mouse lines

TAR4/4 mice were purchased from JAX (stock no.: 012836, B6;SJL-Tg(Thy1-TARDBP)4Singh/J). TDP-43Tg/+ mice were maintained on a B6/SJL hybrid background by crossing with F1 hybrid mice from JAX to propagate the strain (stock no.: 100012, B6SJLF1/J). Mice were housed on a regular light/dark cycle (14:10 h) with ad libitum access to food (LabDiet 5010) and water. All procedures were conducted during the light phase. All protocols for mouse experiments were approved by the Institutional Animal Care and Use Committee and were conducted in accordance with the NIH Guide for the Care and Use of Laboratory Animals

### Intracerebroventricular (ICV) injections

Postnatal day 1 pups were transferred to paper towels over wet ice for hypothermic anesthetization for approximately 5-8 minutes. Pups were confirmed to have been anesthetized by gently squeezing the rear paw and monitoring for the presence of a motor response. A 10ul Neuros Hamilton syringe (Cat.# 7635-01) with needle (32 gauge, point style 4, 0.75 inch length, 30 degree bevel) was used to perform the injection into the cerebral ventricle at 2/5 of the distance directly between lambda and the eye of the pup. The needle was inserted perpendicular to the surface of the skull with the bevel of the Hamilton needle facing toward the midline. Once in place, 3ul of injection solution (15 μg/mL) was slowly dispensed and the needle was held in place for 10 seconds post-injection to prevent backflow before being slowly withdrawn. The pup was then immediately placed onto a warming pad until the normal skin color returned and the pup began moving.

### Phenotype scoring and humane euthanasia end point determination

All phenotype scoring was performed blinded to the genotype or treatment group of the animal, and the order in which the animals were tested each day was random. Gait impairment scoring was used as previously described^3^: score 0 was given for a mouse walking normally: score 1 was given if the mouse had tremor or appeared to limb while walking; score 2 was given if a mouse had severe tremor, severe limp, lowered pelvis or feet pointing away from body during locomotion; score 3 was given if the mouse had difficulty moving forward, with minimal joint movement, feet not being used to generate forward motion, difficulty staying upright, or its abdomen dragging on the ground; score of 4 represented euthanasia end point, where the mouse was unable to right itself within 15 seconds on all 3 out of 3 trials. A humane euthanasia end point was used instead of natural death in survival analysis for the welfare of the animals, but it also served as a more precise end point that was directly dependent on motor dysfunction (an ALS-relevant phenotype) and less dependent on insufficient food and water intake. For tissue collection, all animals were euthanized between 19-32 days of age

### Tissues collected for immunohistochemistry

Mice were euthanized using 2.5% tribromoethanol (0.5 mL/25 g body weight) and transcardially exsanguinated with phosphate-buffered saline (PBS) followed by 4% paraformaldehyde (PFA) perfusion for fixation. Brains were dissected and post-fixed overnight in 4% PFA, then transferred into PBS and shipped to NeuroScience Associates, Knoxville, TN, USA for sectioning and histology.

### Immunohistochemistry in mouse tissue

Brains were treated overnight with 20% glycerol and 2% dimethylsulfoxide to prevent freeze artifacts and multiply embedded into a gelatin matrix using MultiBrain® Technology (NeuroScience Associates, Knoxville, TN). Each MultiBrain block was sectioned coronally at 30 μm. A series of 33 sections, equally spaced at 300 μm intervals throughout the entire hemi brain, were used for staining. IHC staining was performed using anti-NeuN (Millipore, Cat # MAB377), counterstained with Amino cupric silver (de Olmos Stain for degenerating cells).

Whole slide imaging for brain was performed at 200× magnification using the Nanozoomer (Hamamatsu Corp, San Jose, CA, USA) system. 10–12 images per mouse were analyzed using MATLAB (Mathworks, Natick, MA, USA). Tissue sections, neurons, and degeneration stains were detected using color thresholds and morphological operations in MATLAB. Image analysis was performed blinded to experimental grouping and genotype. For brain, 4-5 sections were quantified per region, forebrain (FB) sections quantified fell between (bregma −1.22mm to −2.18) coordinates and the hindbrain (HB) sections quantified fell between (bregma −5.52mm to −6.48) coordinates according to the Allen Brain Atlas. For spinal cord 10 sections from lumbar L3-L5 regions were quantified per animal/stain.

### Primary cultures

#### Mouse primary cortical neurons preparation

Mouse primary cortical neurons were prepared from C57BL6/N mice (Charles River) E15.5 embryos using Papain Dissociation System (MSPP-LK003150, Worthington) and following manufacturer’s instructions. Primary neurons were plated in plating media containing Neurobasal (Gibco 21103049), 1X B27 Supplement (Gibco 17504044), 1X Pen/Strep (Gibco 15070063), 1 X GlutaMAX™ Supplement (Gibco 35050061) and 10% Horse serum (Gibco 26050-088) for 18 hours. Media was then changed with maintaining media containing the same components of plating media without Horse Serum. Half media changes were performed every 7 days.

#### Mouse primary astrocytes preparation

Mouse primary astrocytes were prepared from postnatal day 2 and 3 (P2-P3) C57BL6/N pups (Charles River). Cortices were dissected, washed in HBSS (Gibco 14175095) and triturated in DMEM complete media (made in-house). After trituration cells were plated in T75 flasks and maintained in complete DMEM media 25% FBS for the 7 days. Media was then replaced with complete DMEM 10% FBS. After 10 days in culture, cells were shaken O/N 37C 220rpm to remove microglia and oligodendrocytes, and then trypsinized and replated in complete DMEM 10% FBS.

#### Co-culture preparation

For astrocytes-primary neurons co-cultures, astrocytes were replated after trypsinization in 96 well plates at a density of 50000 cells per well and cultured in DMEM 10% FBS. One week later, media was switched to neuronal maintaining media and 30000 primary neurons were plated on top of the astrocytes. Half media change was performed every 2 days.

### Treatments of primary neurons, astrocytes and co-cultures

#### Antisense oligonucleotides (ASO)

An *Atxn2* antisense oligonucleotide sequence and control non-targeting (NT) ASO sequence were used for the experiments presented in the manuscript and synthesized in house. In primary neurons and co-cultures, ASOs treatment was delivered by gymnosis at a concentration of 10uM starting 2 days after primary neurons plating for TDP-43 overexpression toxicity studies and 5 days after playing for loss of function toxicity studies.

#### siRNA

Horizon Accell Smart Pool Mouse *Idh1, Abca1* and *Hmgcs1* siRNA and were delivered by gymnosis at a concentration of 1uM. siRNA treatment in primary neurons began 2 days after plating. In co-cultures, siRNA delivery began 5 days post neurons plating.

#### Lentiviral transduction

Overexpression of NLS-GFP or hTDP-43-GFP vectors in primary neurons was achieved by lentiviral transduction of cells with the following constructs (Vector Builder) at a concentration of MOI=5: pLV[Exp]-mPGK>NLS-EGFP and pLV[Exp]-mPGK>hTARDBP[NM_007375.3]/3xGGGGS/EGFP. Primary neurons were transduced at DIV5. If double lentiviral expression was required, overexpression of NLS-GFP or hTDP-43-GFP vectors was postponed to DIV7.

Overexpression of BFP-HMGCS1 or BFP vectors in primary astrocytes was achieved by lentiviral transduction of cells with the following constructs (Vector Builder) at a concentration of MOI=5: pLV[Exp]-mPGK>TagBFP2 and pLV[Exp]-TagBFP2-mPGK>mHmgcs1[NM_001291439.1]. Primary astrocytes were transduced two days after replating.

Overexpression of Idh1-Myc-DDK (NM_010497, Origene MR217871L3V) or control (Origene PS100092V) was achieved by lentiviral transduction of primary neurons at a concentration of MOI=5, 5 days after plating.

Loss of function of TDP-43 in primary neurons was achieved by lentiviral transduction of the following vector pLKO.1-CMV-tGFP, containing sequence against TDP-43 GTAGATGTCTTCATTCCCAAA. Scramble shRNA vector pLKO.1-puro containing non-targeting sequence CAACAAGATGAAGAGCACCAA. Lentivirus transduction was performed at MOI=5, 7 days after plating.

### Immunofluorescence for NeuN counting

Primary neurons and primary co-cultures used for TDP-43 overexpression toxicity were fixed 8 days post lentiviral transduction, while for TDP-43 loss of function they were fixed 7 days post lentiviral transduction, based on cell death occurrence. In both cases, cells were fixed in 4% paraformaldehyde (PFA, Electron Microscopy Services #15710) for 20 minutes at room temperature (RT) and washed twice with Phosphate Saline Buffer (PBS). Cells were then blocked in 3% Albumin bovine serum (BSA, EMD Millipore 126609) diluted in PBS for 30 minutes RT. Cells were then permeabilized in 0.2% Triton X-100 in PBS for 10 minutes, washed once in PBS and incubated again in 3% BSA-PBS for 15 minutes. Consequently, primary antibody against NeuN (EMD Millipore #ABN78) was incubated O/N 4C. The day after, cells were washed 3X with PBS and incubated 1h RT with secondary antibody Alexa anti-rabbit 647 (Invitrogen #A21207). Cells were washed 3X with PBS and images were acquired at 10X or 20X magnification with Incucyte S3 (Sartorius) and analyzed with software Incucyte® Base Analysis Software (Sartorius) using the red object counting function. For statistical analysis, each dot in each experiment represents the average of 4 images from one well per replicate.

### Western Blot analysis

Mouse primary co-cultures were directly lysated in NuPAGE™ LDS Sample Buffer containing 0.1M DTT on ice 8 days post transduction with lentiviruses. Samples were boiled 95C for 5 minutes and 20ul of sample was run on NuPAGE™ Bis-Tris Mini Protein Gels, 4–12% (Invitrogen), and consequently transferred on a iBlot™ Transfer Stack, PVDF using the iBlot 2 Western Blot Transfer System, program P0. Membranes were blocked with Intercept (TBS) Blocking Buffer (LICORbio #927-60001) for 30 minutes RT and then blotted primary antibodies diluted in Intercept T20 (TBS) Antibody Diluent (LICORbio #927-75001) O/N 4C. After 3X washing with TBST 0.1% Tween-20, membranes were blotted with IRDye® Secondary Antibodies (LICORbio) for 1h RT. Membranes were washed again 3X with TBST 0.1% Tween-20 and then acquired with Odyssey® DLx (LICORbio) and analyzed with software Empiria Studio 2.1 (LICORbio).

Primary antibodies:

| Protein | Company | Catalogue Number | Specie | Dilution |
| --- | --- | --- | --- | --- |
| HMGCS1 | ProteinTech | 17643-1-AP | Rabbit | 1:1000 |
| GAPDH | Sigma-Aldrich | G8795 | Mouse | 1:10000 |

### RT-qPCR

RNA was extracted from mouse primary cells (8 days post transduction with lentiviruses) and mouse brain tissue using the RNeasy Plus Mini Kit (QIAGEN #74134) following manufacturer’s instructions. After RNA quantification with Nanodrop8000 (Thermo Fisher Scientific), RT-qPCr was performed using 100ng of RNA with TaqMan™ RNA-to-CT™ 1-Step Kit (Applied Biosystems #4392938) according to the manufacturer instructions. RT-qPCR was run on instrument QuantStudio™ 7 Flex (Applied Biosystems). Analysis was performed according to the delta-delta Ct function, normalizing each gene of interest of housekeeping gene *Gapdh*. TaqMan Gene Expression Assays used are the following (Thermo Fisher Scientific): *Ataxin-2* (Mm00485946_m1), *Hmgcs1* (Mm01304569_m1), *Cth* (Mm00461247_m1), *Gapdh* (mm99999915_g1), *Idh1* (Mm00516030_m1), *Hprt1* (Mm03024075_m1).

### Seahorse ATP Rate Assay

Seahorse ATP Rate Assay Kit (Agilent Technologies #103592-100) was used for this experiment according to the manufacturer’s protocol. Primary neurons media was switched to Seahorse XF DMEM Medium, pH 7.4 (Agilent technologies #103575-100) with 10 mM of XF glucose (Agilent technologies #103577-100), 1 mM of XF pyruvate (Agilent technologies #103578-100), 2 mM of XF glutamine (Agilent technologies #103579-100) prior experiment. Cells were plated at a density of 25000 cells per well. Cells were analyzed 8 days post lentiviral transduction. Every experiment is normalized by the number of cells as counted per Hoechst 33342 staining (BD Pharmingen #561908) using the Agilent BioTek Cytation 1 (Agilent Technologies). Experiments were run with the Seahorse XFe96 Analyzer (Agilent Technologies) and analysis was performed with the Seahorse Wave Controller Software 2.6 (Agilent Technologies) exporting the data with the report generator provided with the software.

### Mass Spectrometry-based Proteomics Experiment in Yeast

#### Yeast proteomics sample preparation

Yeast cells were harvested and lysed in HEPES buffer (20mM, pH 8.0) containing 9M urea and phosphatase inhibitors (1mM sodium orthovanadate, 2.5 mM sodium pyrophosphate, and 1.0 mM ß-glycerophosphate). Lysates from 4 experimental conditions were generated for global proteome analysis, resulting in a total of 18 samples. The conditions included: WTwith GFP (n=4), WT with TDP-43 (n=5), *pbp1*Δ with GFP (n=4), and *pbp1*Δ with TDP-43 (n=5). Lysates were sonicated followed by centrifugation at 20,000 g for 20 min at 15 °C. Protein concentration was measured using Bradford assay (BioRad, Hercules, CA). Proteins were reduced with 5 mM dithiothreitol (DTT) at 37 °C for 1h followed by alkylation with 15 mM iodoacetamide (IAA) at room temperature (RT) for 20 min in the dark. Samples were diluted to a final concentration of 2M urea and subjected to sequential digestion: first with Lys-C (Wako, Japan) at 37°C for 2h, followed by trypsin (Promega, Madison, WI) at 37°C overnight, using an enzyme-to-substrate (E:S) ratio of 1:50. The digest mixtures were acidified with 20% trifluoroacetic acid (TFA) prior to solid-phase extraction using C_18_ cartridge (Waters, Milford, MA). Peptides were eluted with 60% acetonitrile (ACN)/0.1% TFA followed by peptide concentration measurement using a quantitative colorimetric peptide assay kit (Thermo, San Jose, CA). Equal amounts from each condition (100 µg) were subjected to lyophilization overnight.

The dried peptide mixtures were reconstituted in 100 µL of HEPES (100 mM, pH 8.5). TMTPro reagent, channels 126-135N (20 µL at 500 µg/50 µL in ACN), was added to each of the 18 samples. Labeling was performed at RT for 1 h. A small portion (2 µL) from each condition was mixed, desalted, and analyzed to determine labeling efficiency. The reaction was quenched with 5 µL of 5% hydroxylamine once labeling efficiency was determined to be at least 95%. Samples were mixed followed by acidification using 20% TFA and lyophilized overnight. The 18-plex TMT labeled peptide mixture was desalted using C_18_ cartridge (Waters, Milford, MA). Peptides were eluted using 60% ACN/0.1% TFA. Sample was lyophilized overnight and subjected to high pH reverse phase fractionation on the Agilent 1100 HPLC system (Agilent Technologies, Santa Clara, CA). Peptide mixture was reconstituted in 75 µL of solvent A (5% ACN/50 mM ammonium bicarbonate, pH 8.0) and separated on a Zorbax 300Extend-C_18_, 3.5 µm, 4.6 x 150 mm column (Agilent Technologies, Santa Clara, CA) at a flow rate of 0.5 mL/min. A gradient from 15-45 % solvent B (90% ACN/50 mM ammonium bicarbonate, pH 8.0) was applied over 49 min with a total run time of 75 min. Ninety-six fractions were collected at 0.63 min interval and every 25^th^ fraction was combined into a set of 24 fractions. Fractions were acidified with 20% TFA, dried and desalted using SDB tips (GL Sciences, Torrance, CA) prior to mass spectrometry analysis.

#### Liquid chromatography mass spectrometry (LC-MS) analysis for yeast

Desalted peptides were reconstituted in 2% ACN/0.1% formic acid (FA)/water and loaded onto Aurora Series 25 cm x 75 um I.D. column (IonOpticks) using a Dionex Ultimate 3000 RSLC nano Proflow system (ThermoFisher). Peptide separation was performed at 300 nL/min with a 2-step linear gradient where solvent B (0.1% FA/2% water/ACN) was increased from 4% to 30% over 70 min then from 30% to 75% B over 4.9 min with a total analysis time of 100 min. Peptides were analyzed using an Orbitrap Eclipse instrument (ThermoFisher Scientific, San Jose, CA). A real-time search against a yeast database was employed using the capabilities built within the Xcalibur method editor. Protein-closeout was employed (3 distinct peptides/protein/run. Precursor ions (MS1) were analyzed in the Orbitrap (250% normalized AGC target, 120,000 mass resolution, 50 ms maximum injection time) with 15 most abundant species were selected for MS2 fragmentation. Ions were filtered based on charge state ≥ 2 (z 2-6) and monoisotopic peak assignment and dynamic exclusion (20 s ± 10 ppm) was enabled. Each precursor was isolated at a mass width of 0.5 Th followed by fragmentation using collision-induced dissociation (CID at 30 NCE), 150% normalized AGC target with a maximum injection time of 100 ms. Multiple fragment ions were isolated using synchronous precursor selection (SPS) prior to MS3 HCD fragmentation (40 NCE, SPS notches = 8, 600% normalized AGC target, maximum injection time of 350 ms). MS3 scans were analyzed in the Orbitrap at 50,000 resolution. Raw datafiles were deposited into the MassIVE repository with the identifier: MSV000099990, reviewer login = MSV000099990_reviewer, password = ATXN2_Jovicic).

#### Data Analysis of DDA-TMT data for yeast

MS/MS data was searched using the Comet search algorithm against a concatenated forward-reverse target-decoy database (UniProtKBconcat, 2021) consisting of yeast proteins and common contaminant sequences. Spectra were assigned using a precursor mass tolerance of 20 ppm and fragment ion settings for low resolution MS/MS with a fragment ion bin tolerance and bin offset of 1.0005 and 0.4, respectively. Static modifications included carbamidomethyl cysteine (+57.0215 Da), TMT tag (+ 304.2071 Da) on both the N-termini of the peptides and lysine residues. Variable modifications included oxidized methionine (+15.994 Da) and TMT tag on tyrosine residues (+304.2071 Da). Trypsin specificity with up to 1 miscleavage was specified. Peptide spectral matches were filtered at 1% false discovery rate (FDR). The TMT reporter ion quantification was performed using an in-house Mojave module^25^ by calculating the highest peak within 20 ppm of theoretical reporter mass windows and correcting for isotope purities.

R package MSstatsTMT v.2.4.1^26^ was used to preprocess PSM-level quantification before statistical analysis, to have protein quantification and to perform differential abundance analysis. MSstatsTMT estimated log2(fold change) and the standard error by linear mixed effect model for each protein. The inference procedure was adjusted by applying an empirical Bayes shrinkage. To test two-sided null hypothesis of no changes in abundance, the model-based test statistics were compared to the Student t-test distribution with the degrees of freedom appropriate for each protein and each dataset. The resulting P values were adjusted to control the FDR with the method by Benjamini–Hochberg.

Gene Ontology (GO) enrichment analysis of yeast dysregulated proteins was performed using DAVID 6.8 (Database for Annotation, Visualization and Integrated Discovery) Bioinformatics Resources^27,28^. Protein categories were assigned based on the functional annotation clustering results from DAVID GO analysis.

### Mass Spectrometry-based Proteomics Experiment in Mouse brain Laser Capture Microdissection (LCM) of cortical layer 5 tissue

Polypropylene microwell chips with a 2.2 mm well diameter were fabricated on polypropylene substrates by Protolabs^4^ (Maple Plain, MN) and used for sample collection and proteomic preparation. LC–MS-grade water, isopropyl alcohol, formic acid (FA), chloroacetamide (CAA), tris(2-carboxyethyl) phosphine hydrochloride (TCEP-HCl), and acetonitrile were purchased from Thermo Fisher Scientific (Waltham, MA). N-dodecyl β-D-maltoside (DDM), dimethyl sulfoxide (DMSO, HPLC grade), sodium chloride (NaCl), and ammonium bicarbonate (ABC) were purchased from Sigma-Aldrich (St. Louis, MO). Trypsin was purchased from Promega (Madison, WI).

Tissue dissection and collection were performed using the PALM MicroBeam system (Carl Zeiss MicroImaging, Munich, Germany), equipped with a RoboStage for high-precision laser micromanipulation and a RoboMover for transferring dissected samples. The system enabled accurate sample collection directly into the microwells of microPOTS chips ^4, 5^. Prior to laser dissection, each microwell was preloaded with 1.5 μL of 50% DMSO aqueous buffer containing 100 mM NaCl and 100 mM ABC. Cortical layer-5 tissue was annotated, dissected, and collected based on anatomical location and cytoarchitecture in coronal sections. The tissue samples were collected into corresponding microwells on the microPOTS chip, with each well serving as a biological replicate (one animal).

After sample collection, the microwell chip was incubated at 60 °C to evaporate DMSO^5^. Subsequently, 2 μL of cell lysis and protein extraction buffer (2 mM TCEP, 5 mM CAA, and 0.1% DDM in 100 ABC buffer) was dispensed into each well using an HP D100 nanoliter liquid dispenser. The chip was incubated at 60 °C for 30 minutes to facilitate protein reduction and alkylation. Enzymatic digestion was then performed by adding 0.5 μL of trypsin solution (80 ng/μL in 100 mM ABC) to each well, followed by incubation at 37 °C for 10 hours. After digestion, peptides were cleaned using Evotip Pure according to the manufacturer’s protocol. The desalted peptides were collected into a 96-well microwell plate, dried with speed vac, and resuspended in 5μL buffer containing 0.1% FA in water.

### LC–MS analysis for mouse brain

Peptide separation was carried out using the Ultimate 3000 RSLCnano system (Thermo Fisher Scientific). Peptides from the well were fully injected into a 20 μL loop and loaded onto a 2 cm-long trapping column (75 μm i.d., 3 μm particle size, 100 Å pore size, C18) using 0.1% FA in water at a flow rate of 3 μL/min for 5 minutes. The sample was then transferred to an IonOpticks Aurora Elite analytical column (250 mm × 75 μm, 1.7 μm particle size) maintained at 50 °C. The separation gradient was as follows: 3–20% Buffer B (acetonitrile with 0.1% FA) over 100 minutes, 20–35% Buffer B over 10 minutes, 35–90% Buffer B over 1 minute, and held at 90% Buffer B for 7 minutes.

Peptides were ionized via electrospray ionization (ESI) and analyzed using an Orbitrap Eclipse mass spectrometer (Thermo Fisher Scientific) coupled with a FAIMS Pro interface. The ion source operated at 1600 V, with the ion transfer tube heated to 300 °C and the S-Lens RF level set to 30. FAIMS compensation voltage was set to −50 V. Data-independent acquisition (DIA) was performed. The DIA method consisted of MS1 precursor acquisitions in the Orbitrap analyzer with a scan range from 400–900 m/z. MS2 fragment acquisition was also acquired with the Orbitrap analyzer with a scan range set to auto where the default charge set to 2. The MS1 spectra were acquired at a resolution of 120K, a normalized AGC target of 250%, and a maximum injection time of 246 ms. The precursors were fragmented at an HCD collision energy of 32%. The isolation window was set to 20 m/z, maximum ion injection time was set to 54 ms and the normalized AGC target was set 3000%. The MS2 fragment spectra were acquired at a resolution of 30K. Raw datafiles were deposited into the MassIVE repository with the identifier: MSV0000999989 (reviewer login = MSV000099989_reviewer & password = atxn2jovicic).

### Data Analysis of DIA data for mouse brain

The Swiss-Prot protein sequence database for Mus musculus (downloaded on 04/17/2024; 17,214 reviewed protein sequences) was used for database searching. DIA-NN Analysis: DIA-NN (v1.8.1) was used for library-free database searching with default parameters. Trypsin/P was specified as the digestion enzyme, allowing for one missed cleavage. Modifications included N-terminal methionine excision and cysteine carbamidomethylation. Peptide lengths were restricted to 7–30 amino acids. A precursor false discovery rate (FDR) of 1% was applied. Additional settings included “--report-lib-info” and the activation of “FASTA digest for library-free search/library generation” and “Deep learning-based spectra, RTs, and IMs prediction” options. Match between runs (MBR) was enabled to improve peptide identification.

DIA-NN (v1.8.1) was employed for library-free database searching using default parameters. The Mus musculus Swiss-Prot protein sequence database (downloaded on April 17, 2024; 17,214 reviewed protein sequences) served as the reference for database searching. Trypsin/P was specified as the digestion enzyme, allowing for one missed cleavage. Modifications included N-terminal methionine excision and cysteine carbamidomethylation. Peptide lengths were restricted to 7–30 amino acids. A precursor false discovery rate (FDR) of 1% was applied. Additional settings included the “--report-lib-info” option and the activation of both “FASTA digest for library-free search/library generation” and “Deep learning-based spectra, RTs, and IMs prediction” options. Match Between Runs (MBR) was enabled to enhance peptide identification. MS1 quantification was utilized for statistical analysis. Filtering was applied based on the following criteria: Lib.PG.Q.value < 0.01, Lib.Q.value < 0.01, PG.Q.value < 0.01, and Q.value < 0.01. Precursor ions exhibiting more than 40% missing values across all 30 runs were systematically excluded from the analysis. Subsequent processing involved equalized median normalization, imputation for missing precursor ions, and protein roll-up, all conducted using MSstats v4.12.0. A linear model was used for statistical analysis, utilizing the lm function from the base R `stats` package, with ‘Group’ and ‘mice’ replicate incorporated as variables. Pairwise comparisons, alongside estimated log2(fold change) and standard error, were determined using the emmeans package. To test the two-sided null hypothesis of no changes in abundance, model-based test statistics were compared against the Student’s t-test distribution, applying degrees of freedom appropriate for each protein and dataset. Resulting p-values were adjusted to control for FDR using the Benjamini-Hochberg method. Pathways analysis identified the pathways from the QIAGEN Ingenuity Pathway Analysis library of canonical pathways that were most significant to the data set. Molecules from the data set that met the *P* adjusted cutoff of 0.01 and were associated with a canonical pathway in the QIAGEN Knowledge Base were considered for the analysis. The significance of the association between the data set and the canonical pathway was measured in two ways: 1) A ratio of the number of molecules from the data set that map to the pathway divided by the total number of molecules that map to the canonical pathway is displayed; and 2) A right-tailed Fisher’s Exact Test was used to calculate a p-value determining the probability that the association between the genes in the dataset and the canonical pathway is explained by chance alone. 3) In many cases a z-score was calculated to indicate the likelihood of activation or inhibition of that pathway.

### Cell-Type Marker Analysis

To address the mixed cellular composition of layer V microdissections, we performed a cell-type marker enrichment analysis using validated markers for major cell types present in cortical tissue. This approach acknowledges that tissue microdissections contain multiple cell types (neurons, glia, and vascular cells) rather than purified populations, and provides evidence for cell-type-specific proteomic signatures without requiring single-cell isolation or flow cytometry.

### Marker Selection and Detection

Cell-type-specific protein marker sets were curated for the following cell types: neurons (13 markers), astrocytes (6 markers), oligodendrocytes (9 markers), oligodendrocyte progenitor cells/OPC (5 markers), microglia (8 markers), endothelial cells (7 markers), and pericytes/smooth muscle cells (10 markers). Detected markers were cross-referenced against our proteomics data to calculate the percentage of each marker set present in the dataset (Supplementary Figure 3C). Markers not detected by mass spectrometry were excluded from downstream analysis to ensure statistical validity.

### Z-Score Normalization and Marker Heatmap

Protein abundance values were standardized across all samples in TAR4/4 Atxn2 ASO and TAR4/4 Ctrl ASO) using row-wise z-score transformation:

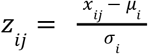

where *x_ij_* is the abundance of protein *i* in sample *j*, *μ_i_* is the mean abundance across all samples, and *σ_i_* is the standard deviation. This normalization accounts for differences in baseline protein expression levels while preserving relative differences between samples and conditions. Detected marker proteins were organized into cell-type groups and visualized as a hierarchical heatmap with z-score values color-coded from −3 (low abundance, navy) to +3 (high abundance, firebrick).

### Declaration of generative AI and AI-assisted technologies in the manuscript preparation process

During the preparation of this work, the authors used Claude (Anthropic) to improve language. The authors reviewed and edited the output as needed and take full responsibility for the content of the published article.

## Supplementary Figures

**Supplementary Figure 1.**
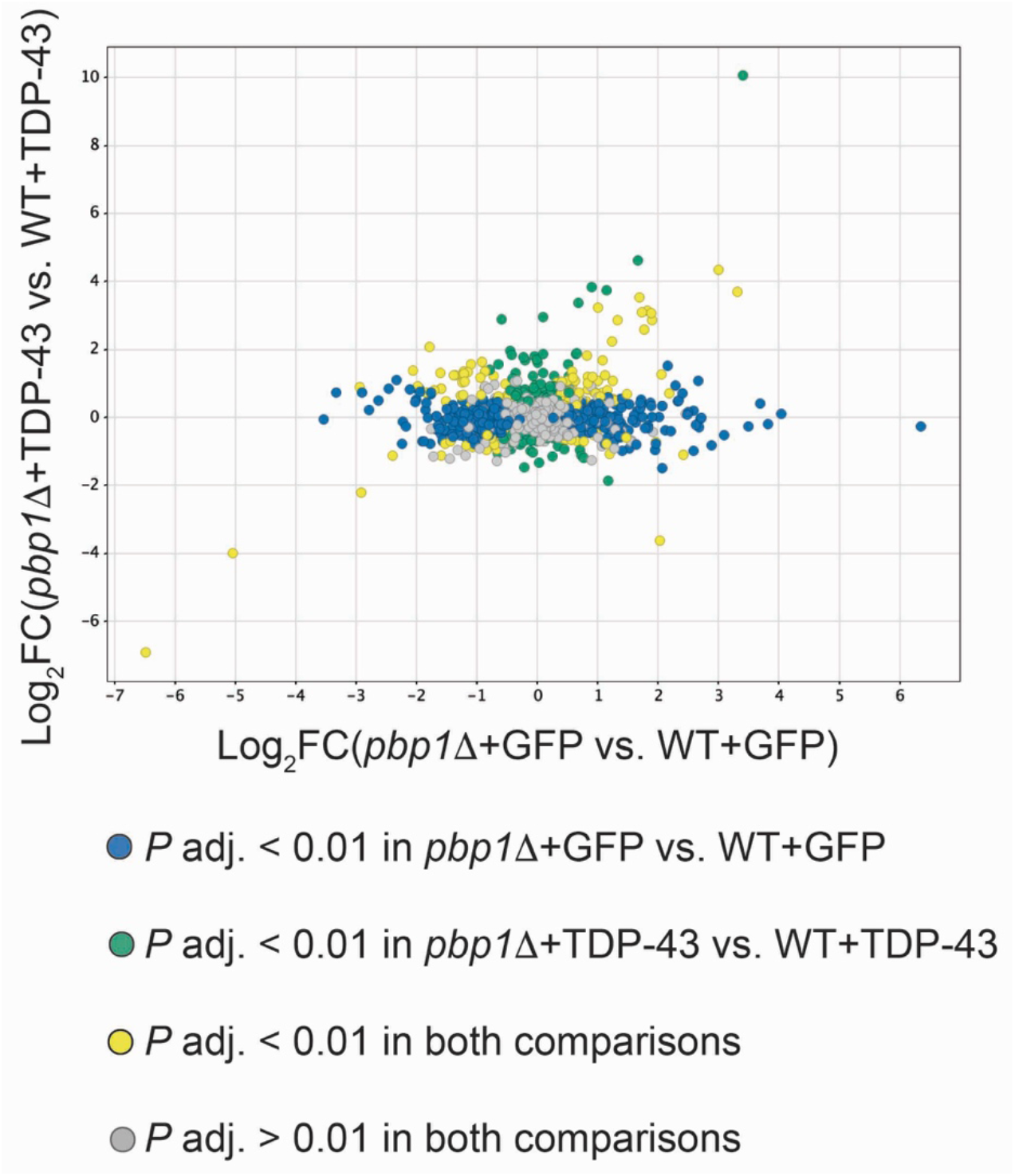
Pbp1D in control GFP-expressing yeast cells results different proteomic response compared to TDP-43-expressing yeast cells (mass spectrometry; P adjusted < 0.01; WT+GFP n=4, WT+TDP-43 n=5, pbp1D +GFP n=4, and pbp1D +TDP-43 n=5).

**Supplementary Figure 2.**
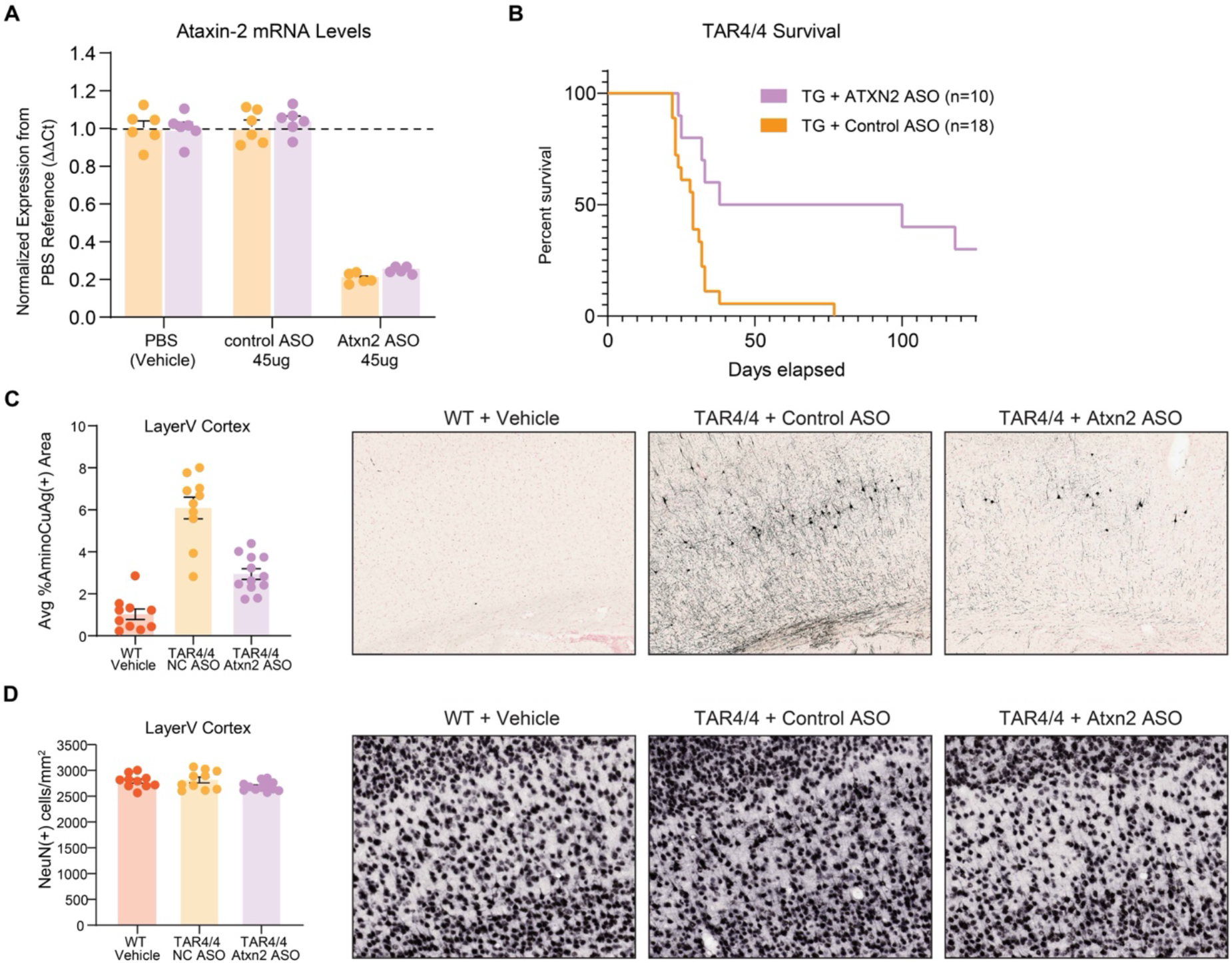
(A) Atxn2-targeting ASO downregulates Atxn2 mRNA in mouse brain (n=6 mice, in each group 3 males and 3 females were used). (B) Atxn2-targeting ASO extends survival of TAR4/4 mice (n=18 mice in the control ASO group, n=10 mice in the Atxn2 ASO group, groups were gender balanced). (C) Atxn2 ASO treatment rescues neurodegeneration in cortical layer 5 of TAR4/4 mice, as assessed by amino-cupric-silver staining (n=10 mice in the WT group, n=10 in the control ASO group, n=10 mice in the Atxn2 ASO group, groups were gender balanced). (D) TAR4/4 mice do not display overt neuronal cell loss in cortical layer 5 (n=10 mice in the WT group, n=10 in the control ASO group, n=10 mice in the Atxn2 ASO group, groups were gender balanced).

**Supplementary Figure 3.**
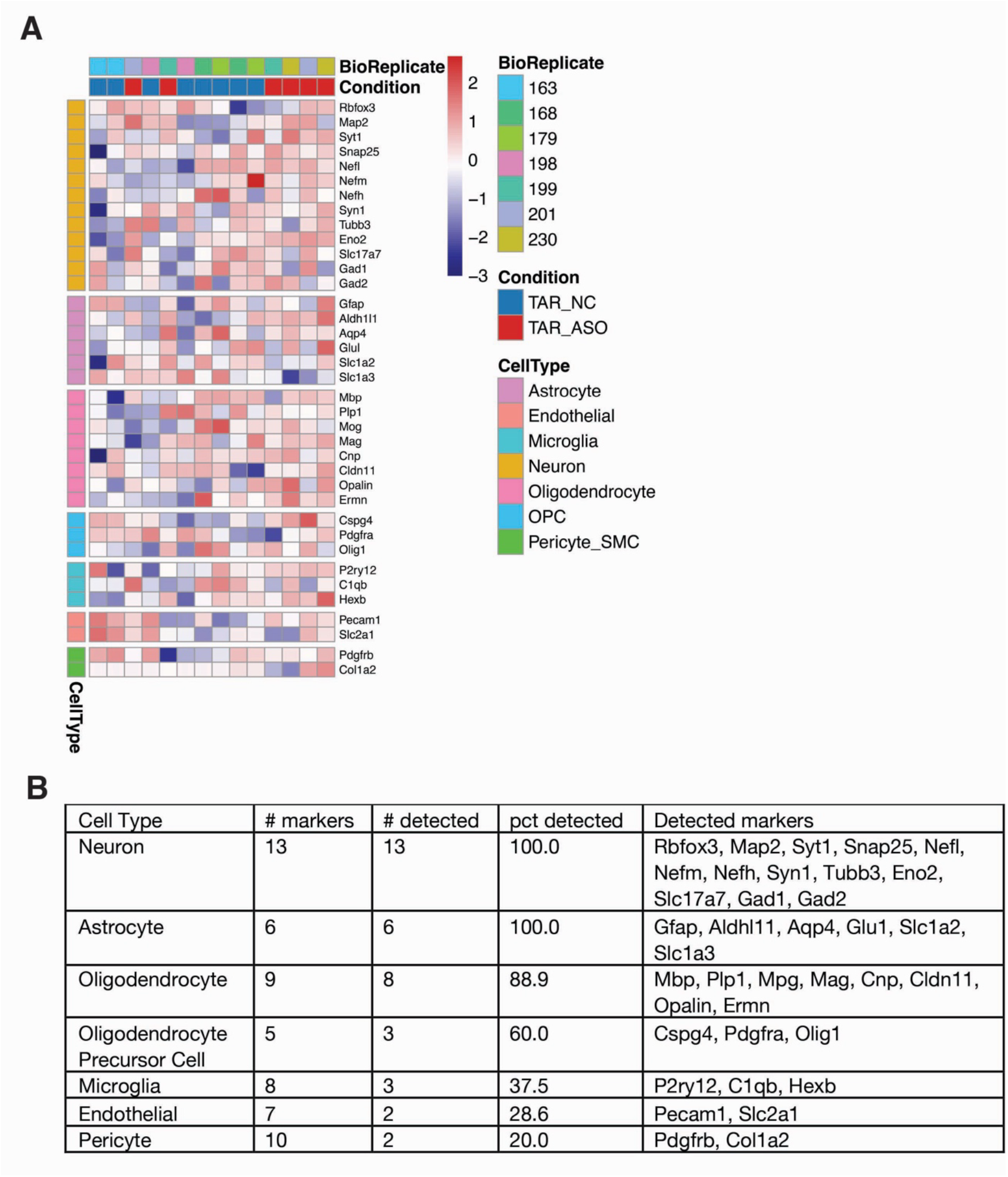
(A) Cell type marker analysis was performed for mass spectrometry data in Figure 1F. The heatmap does not show obvious block-level differences between TAR4/4 mice treated with control or Atxn2 ASO (B) List of cell-type-specific markers used in the cell type marker analysis.

**Supplementary Figure 4.**
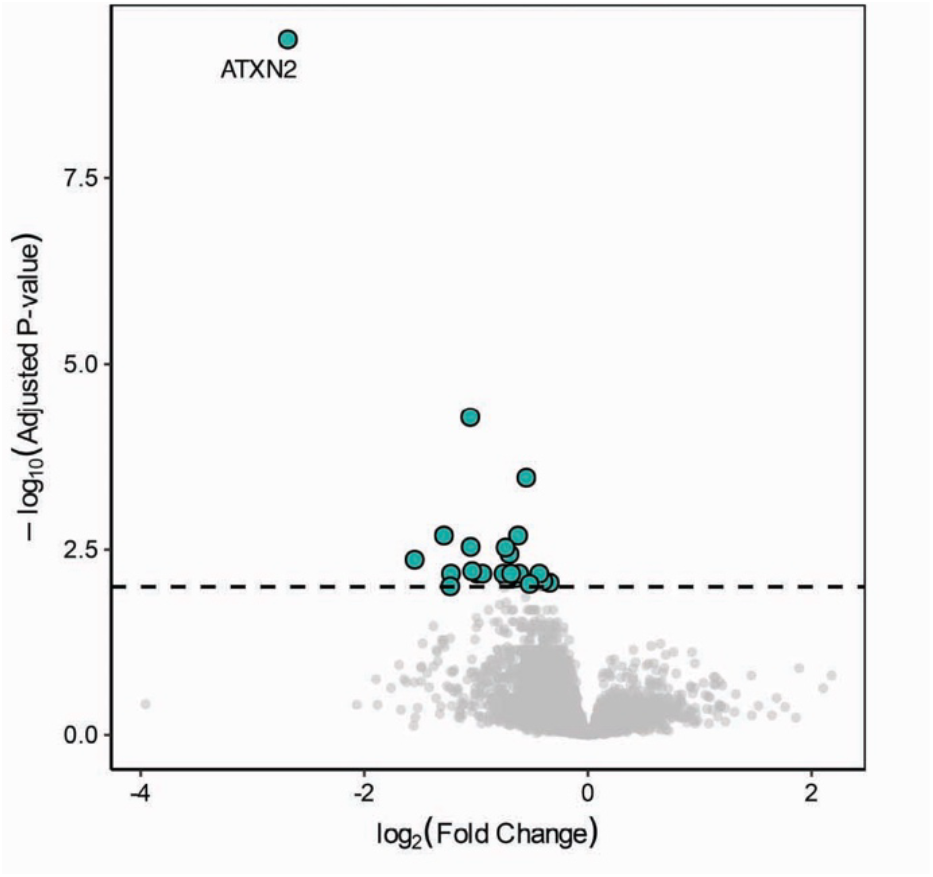
(A) Atxn2 downregulation in wild type mice results in mild downregulation of few proteins. Volcano plot of differentially expressed proteins in layer 5 motor cortex of wild type mice treated with Atxn2 ASO versus control ASO (mass spectrometry; P adjusted < 0.01). n=4 male mice per group.

**Supplementary Figure 5.**
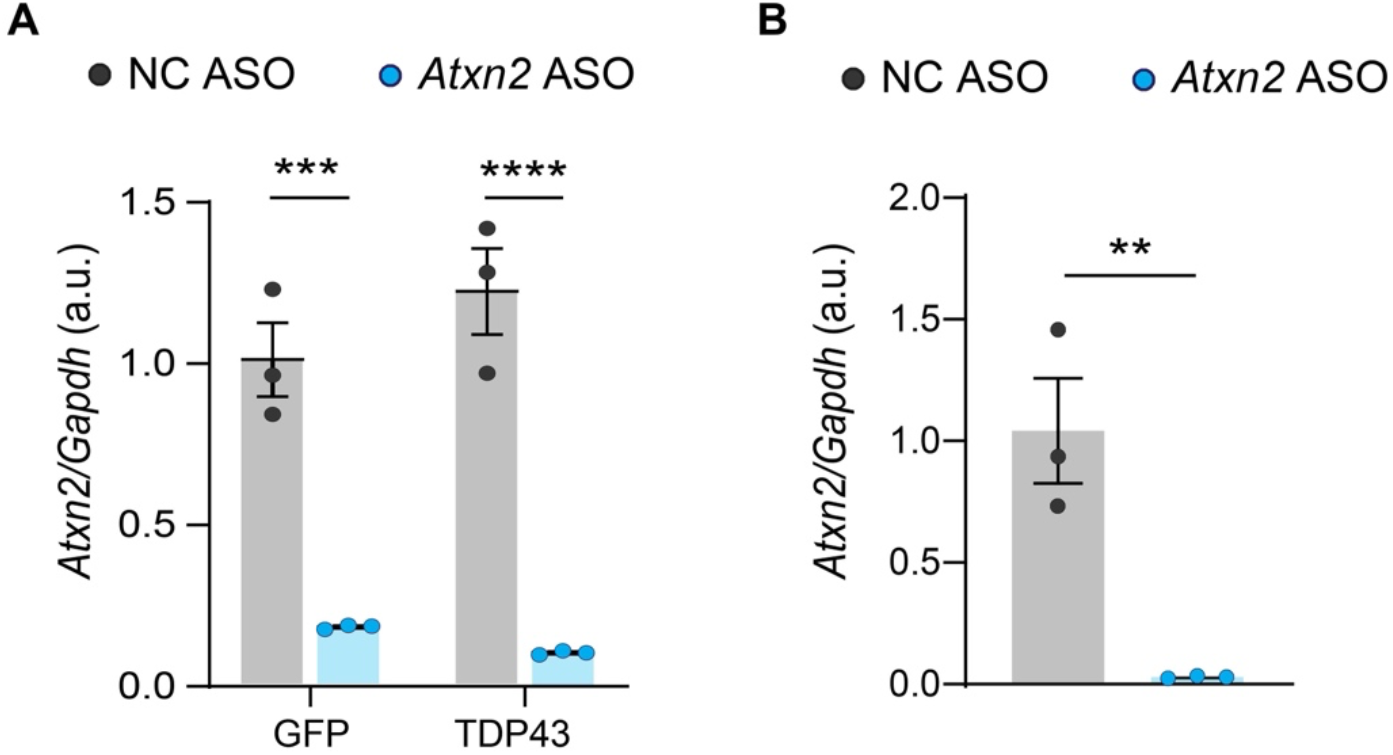
(A) Atxn2-targeting siRNA results in > 80% Atxn2 downregulation in primary neuronal cultures, measured at 72 hours post-treatment. (B) Atxn2-targeting siRNA results in > 90% Atxn2 downregulation in primary neuronal-astrocyte cocultures, measured at 72 hours post-treatment. All expression values were normalized to Gapdh and are presented relative to controls. All data are means ± SEM; statistical significance was assessed by Two-way ANOVA followed by Tukey’s multiple comparison test (in a), or unpaired t-test (b) with * P adjusted < 0.05, ** P adjusted < 0.01, *** P adjusted < 0.001, and **** P adjusted < 0.0001, ns is nonsignificant.

**Supplementary Figure 6.**
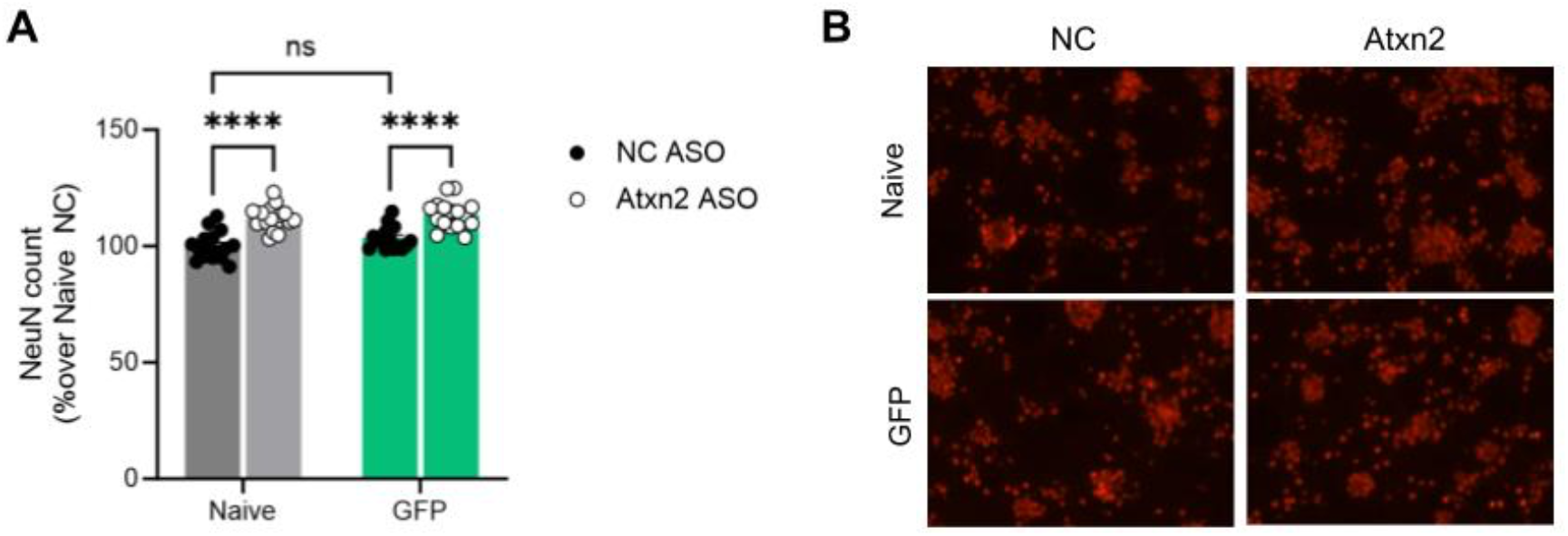
(A) Atxn2 ASO improves survival of non-lentivirus-transduced naïve cultures and GFP expressing cells. GFP expression has no effect on neuronal viability when compared to naïve cells, as assessed by NeuN+ cell counts. All vCellalues are normalized to naïve cells treated with control non-coding (NC) ASO. All data are means ± SEM; statistical significance was assessed unpaired t-test with **** P < 0.0001. (B) Representative images of NeuN images quantified in (A).

**Supplementary Figure 7.**
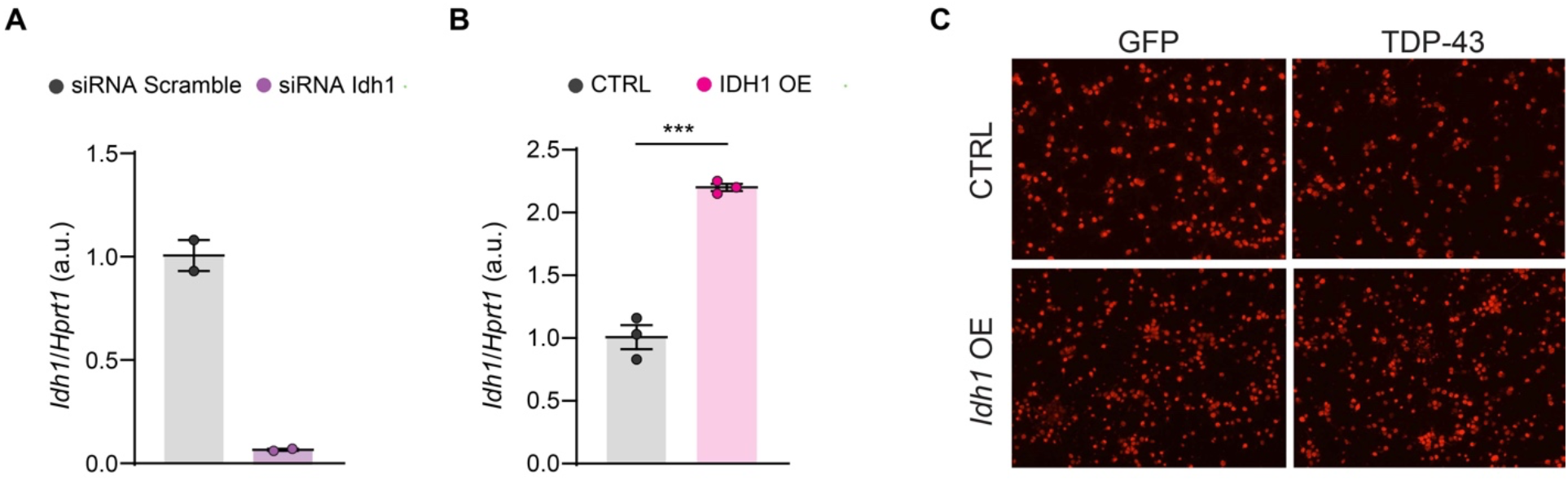
(A) Idh1-targeting siRNA results in > 90% Idh1 downregulation in primary neuronal cultures, measured at 72 hours post-treatment. (B) IDH1 overexpression (OD) results in 2.2-fold protein overexpression at the level of mRNA in primary neuronal culture, measured at 72 hours post-treatment. (C) Representative images of NeuN-labeled neurons quantified in Figure 3G. Expression values in (A) and (B) were normalized to Hprt1 and are presented relative to controls. All data are means ± SEM; statistical significance was assessed unpaired t-test (B) with *** P < 0.001.

**Supplementary Figure 8.**
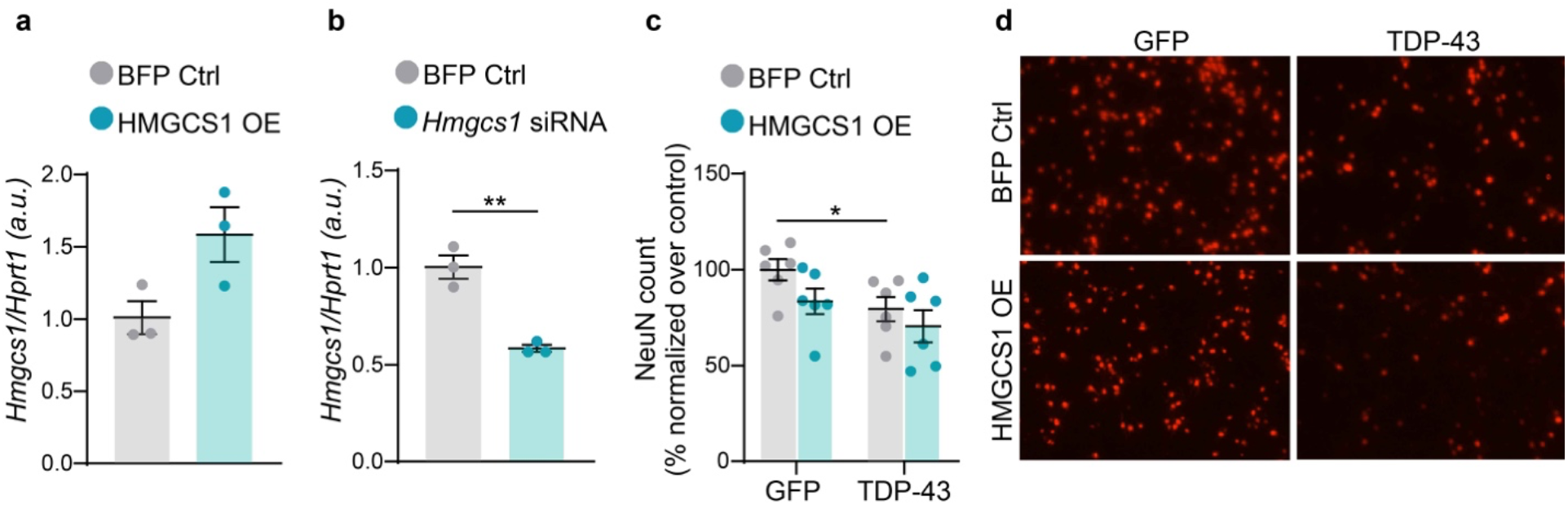
(A) Relative mRNA levels of Hmgcs1 quantified by RT–qPCR in primary astrocytes treated with BFP-control or Hmgcs1 overexpressing lentiviruses. (B) Relative mRNA levels of Hmgcs1 quantified by RT–qPCR in primary astrocytes treated with siRNA Scramble or siRNA against Hmgcs1. In both experiments expression values were normalized to Hprt1 and are presented relative to controls. Data are means ± SEM; statistical significance was assessed by unpaired t-test * P < 0.05. (C) Quantification of primary neuron survival in control (GFP) or TDP-43 overexpressing conditions, overexpressing BFP or HMGCS1. Data are normalized to GFP with BFP condition. (D) Representative images of NeuN-labeled neurons quantified in (C).

